# p21-activated kinase regulates Rab3a vesicles to repair plasma membrane damage caused by Amyloid-β oligomers

**DOI:** 10.1101/2025.02.11.637764

**Authors:** Deepak Kunhi Valappil, Priyadarshini Veerabhadraswamy, Shylaja Partha Sarathi, Prakhyath Hegde, Jeevan M Gowda, Anujith Kumar, Nikhil R Gandasi, Sangeeta Nath

## Abstract

The interaction of amyloid-β (Aβ) peptides with the plasma membrane (PM) is a potential trigger that initiates the formation of higher-order aggregates, membrane alterations/damage, and progressive neurotoxicity in Alzheimer’s disease (AD). In a previous study, we showed that oligomers of Aβ_1-42_ (oAβ_1-42_) induced PM damage, resulting in PM repair cascade via lysosomal exocytosis coupled with endocytosis, and facilitation of tunneling nanotubes (TNTs)-like membrane protrusions to promote direct cell-to-cell transfer of aggregates. In this study, we demonstrated that PM damage induced by oligomers of the aggregation-prone peptide Aβ_1-42_ significantly facilitates PM repair by enhancing phosphorylated p21-activated kinase 1 (pPAK1)-dependent endocytosis and Rab3a-dependent exocytosis in SH-SY5Y and SK-N-SH neuronal cells compared to control and oAβ_1-40_ treated cells. We studied the kinetics of pPAK1-dependent endocytosis and the fusion of EGFP-Rab3a vesicles near the PM using total internal reflection fluorescence (TIRF) microscopy. IPA-3, a selective non-ATP competitive inhibitor of PAK1, inhibits endocytosis of oAβ peptides and Rab3a-dependent PM repair. Further, shRNA-mediated knockdown of the Rab3a gene inhibits pPAK1 and disrupts PM repair. Repair of damaged PM is a vital protective mechanism for non-proliferative cells like neurons, as disruption in PM repair leads to gradual neuronal cell death. However, there was no explicit understanding of PM repair in response to Aβ oligomers. This study revealed the interconnected action of Rab3a and pPAK1 in PM repair in response to oAβ-mediated damage, and its potential correlation in AD pathogenesis.

## Introduction

Neurodegenerative diseases are characterized by the progressive development of pathology. Alzheimer’s disease (AD) is the most prevalent form of dementia among them. The oligomers of amyloid-β (oAβ) are considered potentially neurotoxic. Mutations affecting APP production, processing, and clearance modulate pathogenic Aβ aggregates formation (1). β- and γ-secretases cleave APP at different sites to generate Aβ-peptides with varying lengths. Hydrophobicity of the c-terminal amino acid residues dictates the aggregation propensity of the peptides. The two extra hydrophobic residues (Ile41 and Ala42) make a difference in aggregation propensities between Aβ_1-42_ and Aβ_1-40_. In addition, the membrane environment promotes the transition of monomeric Aβ to the aggregation-prone β-sheet conformation, followed by rapid self-aggregation into pre-fibrillar oligomers and then fibrils (2). A toxic cascade has been triggered by oligomers that cause PM damage, lipid oxidation, and formation of ion-permeable membrane pores (3–5). Toxic aggregates of Aβ bind to lipid membranes, accelerating the formation of progressively more fibrillogenic Aβ aggregates. This, in turn, disrupts the structure and function of the membrane bilayer, leading to neuronal toxicity. After initiation, neuropathological toxicities spread in a ’prion-like’ manner, likely through intercellular transfer of pathogenic aggregates (4, 6, 7).

Aβ peptides originate from the extracellular part of the PM, then they aggregate upon interaction with the membrane lipids in the extracellular space. Endo-lysosomal vesicles then take up the aggregates inside the cells. Several researchers studied the internalization of soluble oAβ via neurons and their fates. Clinical and animal model studies suggest that soluble oAβ, not extracellular deposits, initiates and drives the disease (8). Our previous study showed that neuronal cells internalize oAβ via clathrin-independent endocytosis (CIE) pathway (4). One of the important functions of CIE is to regulate membrane stress, which helps to maintain cellular homeostasis (9). Recent studies have reported that oAβ follows membrane tension-sensitive actin-dependent CIE, which is linked to the Rho-GTPases signalling axis (4, 10). Furthermore, previous studies have shown that tubulin depolymerization and actin polymerization inhibitors can prevent fluid phase-endocytosis in astrocytes and microglia, thereby inhibiting Aβ uptake (11). The p21-activated kinases (PAK) regulate downstream signalling pathways of Rho-GTPases, and activated states of (phosphorylated) pPAKs are described as functional. Involvement of pPAK1 has mostly been reported in membrane tension-sensitive CIE and actin-dependent endocytosis, including fluid phase IL2Rβ endocytosis, macropinocytosis, and the CLIC/GEEC endocytosis (12).

Our previous study demonstrated that neuronal cells initiate PM repair upon oAβ_1-42_-induced PM damage via lysosomal exocytosis coupled with pPAK1-dependent CIE (4). The massive pPAK1-dependent endocytosis during PM repair cascades causes membrane-actin remodulation and the formation of TNT-like membrane protrusions in the oAβ_1-42_ treated neuronal cells (4). Studies showed that pPAKs may facilitate biogenesis of TNT-like structures and cell-to-cell transfer of pathogens (13, 14). TNTs are nano-sized membrane tubes that connect distant cells through open-ended channels and facilitate direct cell-to-cell transfer of neurodegenerative aggregates, including the Aβ_1-42_ oligomers between distant neurons (4, 15, 16). Cell-to-cell transfer possibly promotes the spread of pathology in neurodegenerative diseases.

Several *in vitro* studies on artificial lipid bilayers showed that the interaction of lipids with oAβ caused membrane damage. Aggregates of Aβ binding to the PM increase membrane permeability and leakage (3, 6). However, only a few (4, 5) studies showed neuronal membrane repair upon oAβ induced PM damage and how defects in PM repair contribute to neurotoxicity (17).

Studies have shown that two members of the Rab family (Rab3a and Rab10) of small GTPases are essential for lysosomal exocytosis-mediated resealing of the PM after damage by mechanical shear force or pore-forming toxin (18). Rab3a silencing disrupts lysosome positioning and ends up localizing lysosomal vesicles to the perinuclear region, resulting in the subsequent inhibition of PM repair. The non-muscle myosin heavy chain IIA (NMHC IIA) identified as part of the lysosome exocytosis complex (that includes Slp4-a and Rab3a) manages the positioning of lysosomes at the periphery of the cell membrane (19). Study has shown that oAβ_1-42_ can directly insert into and damage the PM, inducing a PM repair process similar to that caused by pore-forming bacterial toxins, streptolysin-O (SLO) (5). A recent study shows that elevated levels of Aβ lead to dysfunctional PM repair by suppressing dysferlin, a crucial protein involved in membrane trafficking and vesicle fusion within the repair patch (17). The defective PM repair process is exorbitant for terminally differentiated neurons, as it can lead to neurodegeneration. Thus, understanding the mechanism of PM repair at the molecular level in response to damage induced by toxic oligomeric Aβ peptides is highly important.

In this study, we have observed that pPAK1-dependent internalization of oAβ peptides modulates Rab3-dependent vesicle fusion/exocytosis to repair oAβ induced PM damage. We were able to show for the first time, pPAK1-mediated endocytosis and Rab3a-dependent vesicle fusion at the PM in response to two different types of oAβ peptides, the aggregate-prone oAβ_1-42_ and the normal physiological oAβ_1-40_. We used a total internal reflection fluorescence (TIRF) microscope to examine the dynamics of these processes. We observed oAβ_1-42_ treatment-induced PM damage causes faster PM repair response via Rab3a vesicle fusion at PM. The rapid PM repair response increases the formation of more TNT-like membrane protrusions between distant cells, likely due to impaired membrane recycling homeostasis. IPA-3, a selective non-ATP competitive inhibitor of PAK1, inhibits the internalization of oAβ and subsequent PM repair. Inhibition in Rab3a expression using shRNA disrupts PM repair, which leads to neurotoxicity. Thus, the study is important and relevant, and it will open up novel avenues for therapeutics by targeting molecules related to PM repair pathways in AD.

## Experimental Procedures

### Cell culture

SH-SY5Y (ECACC: catalogue #94030304) and SK-N-SH (NCCS, Pune, India, repository) human neuroblastoma cells were grown in DMEM/F12 1:1 (Gibco) supplemented with 15% FBS (Corning), 1% Glutamax (Gibco), 1% Non-essential amino acids (Gibco) and 1% PSN (Penicillin-Streptomycin-Neomycin Mixture, Thermo Fisher Scientific). The cells were plated with 0.05×10^6^ cells in 100 mm dishes (Corning), 0.03×10^6^ cells in 24 well plates (Corning), and 0.005 x10^6^ cells in 14 mm diameter glass-bottom dishes, made of # 1.5 coverslip (Cellvis, D35-14-1.5-N). Experiments were performed in both the SH-SY5Y and SK-N-SH cell lines; results were similar in both cell lines. In the manuscript, we have presented the representative results from SH-SY5Y cells. Morphologically, the two cell lines are the same. Their similarities in terms of morphology are shown in (Figure S1).

HEK293T cells were cultured in MEM(E) (Merck) with 10% FBS (Corning) and 1x PSN (Thermo Fisher Scientific). The cells were seeded with 0.05 x 10^6^ cells in 100 mm dishes (Corning).

### Preparation of soluble oAβ

Labelled and unlabelled Aβ peptides (Aβ_1–40_ and Aβ_1–42_) purchased from AnaSpec, were reconstituted in HFIP (1,1,1,3,3,3-hexafluoro-2-propanol) (TCI) and subsequently lyophilized into 0.05 mg peptide aliquots. Following this, the lyophilized Aβ peptides were rehydrated to a concentration of 5 mM in Me2SO (Molecular grade, Himedia), and then further diluted to a concentration of 100 μM in DMEM without phenol red (Gibco) at pH 7.4. The solution underwent immediate vortexing and sonication for 2 minutes, followed by a 24-hour incubation at 4°C (4). Before the experiments, the oAβ peptides were characterized, as detailed in our previous publications (20, 21), using transmission electron microscopy (TEM) imaging. The oligomers of Aβ peptides were resuspended in 1X PBS, on a carbon grid, coated with uranyl acetate. The resuspended oligomers were dried in a desiccator and then visualized using TEM microscope FEI Tecnai T20 at 200 KV.

### Treatment with oAβ peptides

Upon reaching 70% confluency, the cells were incubated with prepared oAβ, directly from a 100 μM stock (dissolved in DMEM without phenol red) and was added to the medium without FBS supplement to achieve a final concentration of 1 μM. The treated samples were then incubated for different periods as per the experimental conditions at 37°C.

### Intracellular localization of oAβ

To investigate whether oAβ peptides were internalized within lysotracker-positive vesicles, fluorescently labelled oAβ-TMR (1μM) (AnaSpec) peptides were introduced into the cells and incubated for 1 hour. Lysotracker was added following a quick wash with 1xPBS and incubated for 5-10 minutes. The cells were fixed with 4% PFA (paraformaldehyde, Sigma Aldrich) for 15 minutes at room temperature and mounted with DABCO dissolved in 90% glycerol before capturing confocal images.

### Immunocytochemistry

Differential immunostainings were performed to gain insights into actin modulation following membrane damage and the distribution of Rab3a for PM repair (22, 23), using pPAK1[phospho-PAK1] (Thr423)/PAK2 (Thr402) antibody (Cell Signaling Technology; CST #2601); f-actin binding phallotoxin phalloidin (A30106, Invitrogen); EEA1 [E4156, Sigmaaldrich]; Lamp1 (PE Conjugated, 2293862, Invitrogen) and Rab3a (PA1-770, Invitrogen).

The control and oAβ (1μM) treated cells were washed with 1X PBS before fixation. The cells were then fixed by adding Karnovsky’s fixative solution for 45 minutes at room temperature. The solution was prepared by using 2% formalin fixative (Sigma Aldrich, Neutral buffered 10%, HT5014-120ML) and 2.5% glutaraldehyde (Sigma, Grade I, 25% in H_2_O) dissolved in 0.1 M phosphate buffer of pH=7.2. After fixation, the primary antibodies were used at a dilution of 1:250 in the incubation buffer and incubated overnight at 4^0^C in a dark moist chamber. The incubation buffer (20X) was prepared by dissolving 0.1 g of saponin (Merck, 8047-15-2) in 5 ml of FBS, then diluting the solution (1X) to 100 ml in PBS. Next day after 24 hrs of incubation, cells were washed with the incubation buffer, then secondary antibodies Alexa flour 488 (Invitrogen (A11070), Goat Anti-rabbit IgG (H+L), 1:1000 dilution), and Lamp1 (PE-conjugated) and Phalloidin iFlour 555 (1:300 dilution) were added incubated in a dark moist chamber for 1.5 hours at room temperature. The cells were mounted with a mounting reagent (Invitrogen) before proceeding to confocal microscopy imaging.

### Western blotting (WB)

Cell lysates were prepared by dissolving cell pellets in RIPA buffer freshly supplemented with 2% PMSF (from a 100mM PMSF stock dissolved in isopropanol) and 2% protease inhibitor cocktail (Sigma Aldrich). The concentration of each unknown lysate sample was determined through the Bradford assay, using BSA as standard. SDS-PAGE (10%) was performed using the cell lysates after mixing with 5x Laemmli buffer (ML121, Himedia) and heated for 5 minutes at 95°C. Proteins from the SDS-PAGE were transferred to a PVDF Immobilon-P membrane (Millipore). Next, the membrane underwent overnight incubation at 4°C with constant agitation, utilizing antibodies against pPAK1 [Cell Signaling Technology, #2601, (1:750)], Rab3a [PA1-770, Invitrogen (1:750)], Lamp1 [611042, BD transduction Lab, (1:750)], and EEA1 [E4156, Sigmaaldrich] (1;750), with loading control antibodies directed at actin (PAS097Hu01, Cloud Clone) or GAPDH (CAB932Hu22, Cloud Clone) in 1:1000. After the incubation with an HRP-conjugated secondary antibody (1:1500; Invitrogen), the ECL kit from Thermo Supersignal was used to develop the WB bands. The density of each band from the ECL images was analysed using Fiji (Fiji is just image J) software.

### Propidium iodide internalization to detect PM damage

Propidium iodide (Sigma Aldrich) uptake serves as an indicator of PM damage. The repair of PM damage can occur through Ca^2+^-dependent lysosomal exocytosis, as previously detailed. Cells were exposed to oAβ (1μM) in DPBS buffer (PBS with Ca^2+^ and Mg^2+^ chloride) at 37°C. DPBS buffer supplemented without or with 5 mM EGTA to maintain the presence and absence of Ca^2+^ respectively (24). Following oAβ treatment (15-30 minutes for live cell imaging), the extent of membrane damage was analyzed by dual nuclear staining with Propidium iodide (PI) and Hoechst. The cells were rapidly washed with media, followed by staining with a PI staining solution (prepared with 1μg/mL of PI stock [1mg/mL in 1xPBS] and 10 μg/mL RNase A [Sigma Aldrich] dissolved in 1xPBS), diluted with 1x PBS (1:100) for 5 minutes at room temperature. After a quick wash with media, the cells were further stained with Hoechst, diluted in 1xPBS (1:2000), for 5 minutes at room temperature. Imaging was conducted using a fluorescence microscope (Olympus IX73) immediately after a rapid media wash and adding fresh media.

### Lentiviral-based shRNA-mediated knockdown

When HEK293T cells reached 60-70% confluency, the transfer plasmid (shRab3a integrated into the pLKO.1 vector, SHCLNG TRCN0000047953, Sigma Aldrich) was transfected alongside packaging plasmids: pMD2.G (addgene id: #12259) and psPAX2 (addgene id: #12260), using FuGENE (Promega) dissolved in OptiMEM (Gibco). In a FACS tube (BD life sciences), 600 μL OptiMEM was mixed with 30 μL Fugene and incubated for 5 minutes at room temperature. Plasmids were isolated using the Qiagen Plasmid Midi kit following the manufacturer’s protocol. The three plasmids (each with a concentration of 4μg-10 μg) were combined in the tube cap. The FACS tube was closed, and the mixture was combined with the Fugene mixture and then incubated for 20 minutes at room temperature. An empty pLKO.1 vector was also mixed similarly to serve as a control. The transfection mixture was added to the HEK293T cells. Viral particles were concentrated by centrifugation at 1500 rpm for 5-10 minutes using centrifugal filters after 48 and 72 hours of transfection. The collected viral particles were then transduced into recipient cells, SH-SY5Y, in DMEM/F12 media containing 0.5% puromycin (Gibco). The cells were passaged for subsequent experiments, and the knockdown efficiency remained consistent up to the 5th passage.

### Cell viability assay MTT assay

The viability of cells was assessed in triplicates using the MTT (3-(4,5-dimethylthiazol-2-yl)-2,5-diphenyl tetrazolium bromide) [HiMedia] reagent. Cells were treated with oAβ (1 μM) at various incubation times, both with and without pre-treatment of cells with IPA-3 (20 μM) for 30 minutes at 37°C. The cells were then incubated with MTT reagent for 2 hours at 37°C. Following this, the media were removed, and the Me2SO [HiMedia]-solubilized Formazan was measured. Formazan (bright orange), the MTT’s reduced product, was measured at 570 nm using an Ensight^TM^ multimode plate reader (PerkinElmer).

### TUNEL assay

The terminal deoxynucleotidyl transferase-mediated dUTP nick end labelling (TUNEL) assay was performed to measure the early and late stages of apoptosis. The TUNEL assay detects DNA breakage by incorporating BrdUTP to the free 3ʹ-hydroxyl termini. An Alexa Fluor 488-labelled anti-BrdU monoclonal antibody is used to detect BrdU. Cells received oAβ (1 μM) treatment at different incubation times, with or without prior exposure to IPA-3 (10 and 20 μM) for 30 minutes at 37°C. APO-BRDU TUNEL assay kit (Thermo Fisher Scientific) was used on the cells to detect the apoptosis cells. Fluorescence microscopy images were used to identify BrdU fluorescence and nuclear DAPI staining using 488 and 405 filters within the nuclei of apoptotic cells.

### Identification and characterization of TNTs

Confocal images saved in Carl Zeiss Image Data format (.czi files) were opened in Fiji. Characterization of TNTs is challenging because there are no specific markers available. The current characterization method involves identifying f-actin-positive hovering membrane channels that maintain connectivity between neighboring cells, even in the fixed cells. The TNTs were identified or distinctly characterized from developing neurons, and these structures were manually counted, as described earlier (25).

### Tracking of EGFP-Rab3a dynamics using TIRF microscope

#### Cells

Cells were seeded on 22-mm gelatin coated coverslips in 6 well plates. Transient transfection with 0.5 µg of EGFP-Rab3a construct was performed using jetPRIME (Polyplus, 101000015). The reaction was terminated after 8 hours, and cells were imaged by maintaining in the extracellular buffer after 36-48 hours of transfection. Extracellular buffer contained 138 mM sodium chloride (NaCl), 5.6 mM potassium chloride (KCl), 1.2 mM magnesium chloride (MgCl_2_), 2.6 mM calcium chloride (CaCl_2_), 17.5 mM D-glucose, and 5 mM HEPES (pH 7.4). oAβ peptides (1 μM) were added to the buffer and incubated for 15-20 minutes before imaging. Control images were acquired before adding oAβ peptides (oAβ_1-40_ and oAβ_1-42_).

#### TIRF-Microscopy

To study Rab3a dynamics, cells were imaged at 37 °C using a TIRF microscope, Nikon ECLIPSE Ti2. Time-lapse images were captured before and after adding oAβ peptides with a 100X/1.45 objective and 488 excitation filter for 5 minutes with 5-second intervals by an ORCA-Flash4.0 V3 digital CMOS camera (Hamamatsu). 16-bit images were acquired with 0.065 µm/pixel resolution.

### Analysis of EGFP-Rab3a dynamics

The density of EGFP-Rab3a puncta was calculated in ImageJ (http://rsbweb.nih.gov/ij) using built-in script called ‘find maxima’ for spot detection (26, 27). The count was then normalized to the area. Rab3a puncta in the TIRF field were marked as ROI (Region of interest) using the image analysis software MetaMorph (Molecular Devices, Sunnyvale, CA, USA). An algorithm of MetaMorph journal was used to obtain the change in the fluorescence intensity over time within the ROI. The average pixel fluorescence of the granule in 1) a 3 pixels diameter central circle (c), and 2) a 5 pixels outer diameter surrounding annulus (a), was read by the algorithm to obtain granule fluorescence ΔF, by subtracting the circle (c) with the annulus value (a) (ΔF=c-a). This ΔF value was considered as per-pixel average for the entire 3 pixels circle.

ΔF values of the puncta were plotted against time (s) (27). The residence time of the puncta was obtained by calculating the time difference between the appearance and disappearance of the puncta within the ROI. Rab3a puncta movement was analyzed using the trackmate function in ImageJ software with a 7-pixel (0.455 μm) blob diameter. The average distance and total distance were calculated as illustrated in Figure 3L-O (28).

### Endocytic vesicles tracking

FITC-conjugated dextran (Sigma; MW 70 kDa) 1 mg/ml was used as an endocytic marker. The endocytosis of oAβ-TMR peptides (1 μM) in SH-SY5Y neuronal cells was observed using a TIRF microscope following the introduction of FITC-conjugated 70 kDa dextran, which serves as a marker for fluid phase endocytosis. The dynamics of endocytic vesicle puncta were observed at the PM in the TIRF plane after incubating the cells with FITC-dextran, along with oAβ-TMR peptides (1μM) for 10-15 minutes. The measurements were also taken with or without prior exposure to IPA-3 (10 and 20 μM) for 30 minutes at 37°C. The endocytosis events were recorded near the PM for 5 minutes with 5 seconds interval using TIRF microscope.

### Analysis of endocytic events

The density of oAβ-TMR and FITC-dextran puncta was calculated using image analysis software MetaMorph. Then, endocytic vesicle puncta in the TIRF field were marked as ROI using MetaMorph. The colocalization of oAβ-TMR and FITC-dextran puncta were analyzed using MetaMorph software. ROIs were marked manually in green channel after identifying the FITC-dextran puncta. These ROIs were transferred to the other channel and the centering was marked by a yes/no choice. ROI positioning within one pixel at the centre of the previously identified oAβ-TMR ROIs was identified and plotted as percentage of colocalization.

### Confocal Microscopy

Immunostained cells were imaged using the Zeiss LSM880 confocal laser scanning microscope (Carl Zeiss, Germany). The pictures were captured using a Plan-Apochromat 40x/1.40 Oil Dic M27 objective, equipped with DAPI, FITC, and TRITC filter sets (Carl Zeiss, Germany). Sequential imaging of the three channels—DAPI, FITC, and TRITC—was performed using lasers with wavelengths of 405 nm, 488 nm, and 561 nm, respectively, and captured with a pixel dwell of 1.02 µs. To observe the cell boundary DIC (differential interference contrast) channel images were sequentially captured along with fluorescence channels. A stack of images was taken with dimensions of x: 224.92 µm and y: 224.92 µm, featuring each pixel at a size of 220 nm² and a z-scaling of 415 nm. Multiple z-stacks were captured, covering the bottom to the top of the cells.

#### Colocalization analysis

The confocal images were used to analyze the colocalization percentage. The expression of two proteins within the same cells were assessed using a color threshold tool in Fiji. The percentage of colocalization or internalization was then determined based on the colocalized area using this tool.

### Statistical Analysis

Statistical analyses were performed using GraphPad Prism 8. To confirm the significance of the analyzed data, one-way ANOVA tests, as well as Student t-tests, were performed according to the experimental parameters specified in the figure legends. The data represent a minimum of n = 3 biological repeats.

## Results

### oAβ peptides internalize to endo-lysosome vesicles via pPAK1-dependent endocytosis

The toxic oAβ peptides can cause damage to the cell membrane of neuronal cells. This damage, in turn, results in the acceleration of endocytosis of these aggregates (4). Here, we have demonstrated that the extent of endocytosis correlates with the aggregation propensity of Aβ-peptides. It is known that Aβ_1-42_ has a high propensity to induce a higher order of toxic oligomers than Aβ_1-40_. Aβ_1-40_ is more relevant in terms of its physiological role and its abundance (2). Our observation shows that Aβ_1-42_ creates larger oligomers as compared to Aβ_1-40_ (Figure 1A-B). The oligomers were characterized by TEM. We observed that fluorescent dye tetramethyl-rhodamine labelled (TMR-labelled) oAβ_1-40_ and oAβ_1-42_ both internalize to lysotracker-positive vesicles of endo-lysosomes in the SH-SY5Y neuronal cells (Figure 1C-D). However, the analysis shows that oAβ_1-42_ colocalize at a significantly higher extent in the lysotracker-positive vesicles after 2 hours of exposure compared to oAβ_1-40_ (Figure 1E). Image analysis shows that lysosomal vesicles containing oAβ_1-42_ accumulation are significantly larger than those containing oAβ_1-40_ (Figure 1F).

**Figure 1:**
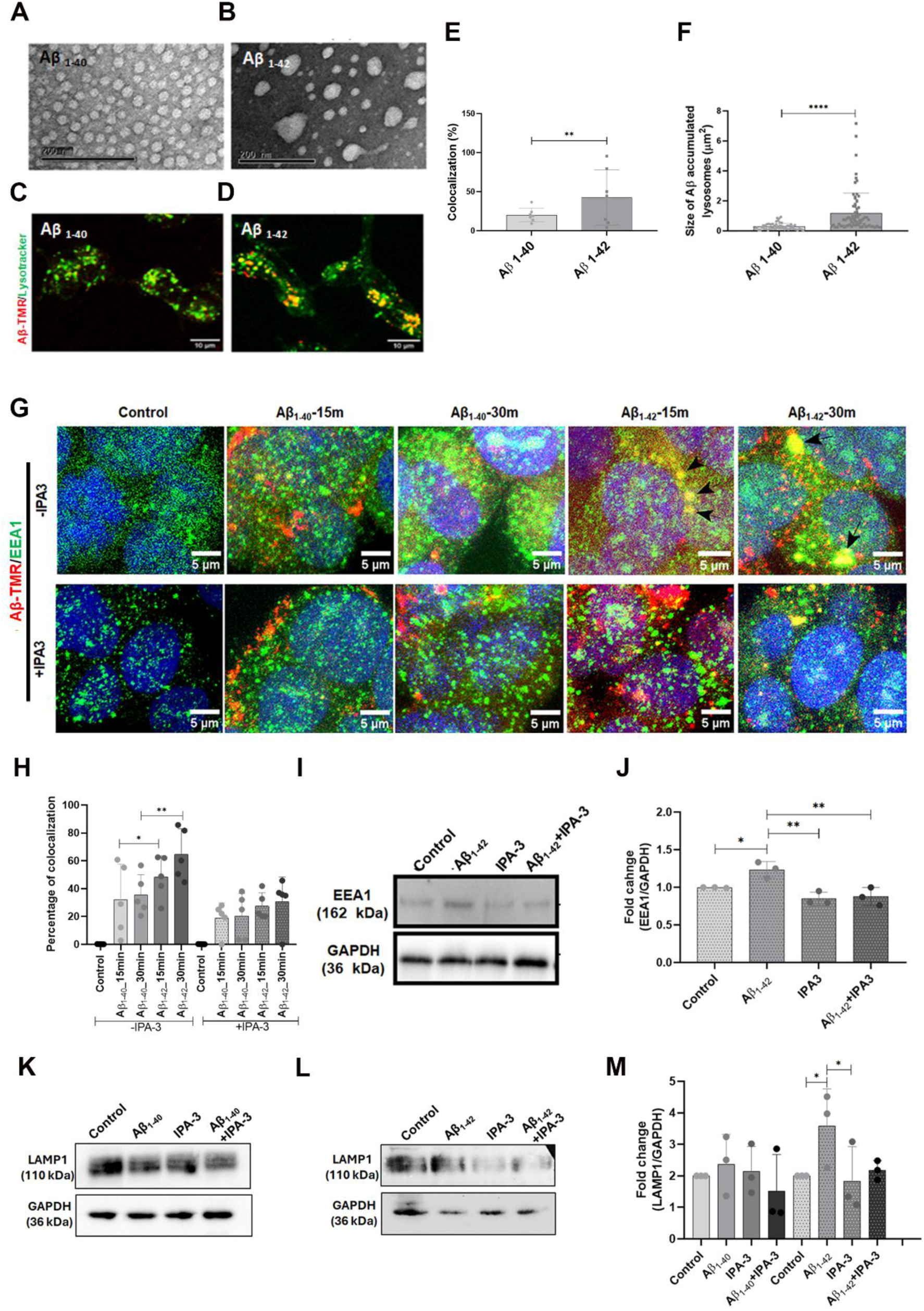
Internalization of oAβ peptides in the endolysosomal pathways in SH-SY5Y cells. TEM images of **A)** oAβ_1-40_ and **B)** oAβ_1-42_. **C, D)** Fluorescently (TMR, red) labelled **C)** oAβ_1-40_ and **D)** oAβ_1-42_ end up to lysotracker positive vesicles (green). **E)** Percentage of colocalization of TMR-labelled oAβ peptides with lysotracker positive vesicles after 2 hours of exposure, and **F)** Size of oAβ accumulated lysotracker positive endolysosomes. **G)** Internalization of oAβ_1-40_-TMR and oAβ_1-42_-TMR into the EEA1 positive early endosomal vesicles. **H)** Graphical representation of oAβ peptides colocalization with EEA1. **I)** WBs of EEA1 Aβ_1-42_ treated cells with and without IPA-3. **J)** Quantification of EEA1 expression. **K,L)** Lamp1in **K)** Aβ_1-40_ and **L)** Aβ_1-42_ treated cells, with and without IPA-3, a specific inhibitor of pPAK1. **M)** Quantification of Lamp1 levels. Data are expressed as mean ± SD, *** p ≤ 0.001. Statistical significance was validated using the student-t test for E and F, and one-way ANOVA for I and K. n=3.

This difference in lysosomal accumulation may arise from oAβ_1-40_ being more degradable or having a lower endocytosis rate. We analyzed the endocytic events of oAβ_1-40_-TMR and oAβ_1-42_-TMR by observing fluorescence puncta near the PM with a TIRF microscope. The result shows that the number of oAβ_1-40_-TMR puncta near the PM is significantly lower compared to oAβ_1-42_-TMR (Figure S2A-B). FITC-conjugated dextran was used as an endocytic marker, as it is known as a fluid phase endocytic marker that involves pPAKs, including IL2Rβ endocytosis, macropinocytosis, and the CLIC/GEEC endocytosis. Our earlier study discovered that oAβ_1-42_ follows a pPAK1-dependent endocytosis and noticed that a small amount of oAβ_1-42_ colocalizes with 70 kDa dextran-FITC when we examined it using a confocal microscope (4). Therefore, we employed FITC-dextran as an endocytic marker and noted a greater degree of partial colocalization between oAβ_1-42_-TMR and dextran compared to oAβ_1-40_-TMR and the control (Figure S2C). Inhibition of endocytosis using IPA-3, a reversible specific inhibitor of PAK1’s autoregulatory domain, results in a decrease in endocytic events (Figure S2D-E) and reduces colocalization with dextran for both oAβ_1-40_-TMR and oAβ_1-42_-TMR (Figure S2F). However, the endocytosis rate of oAβ_1-40_-TMR is significantly slow, making the inhibitory effect of IPA-3 less apparent for the peptide oAβ_1-40_.

Further, the internalization of fluorescently labelled (TMR-labelled) oligomers of Aβ_1-40_ and Aβ_1-42_ within the EEA1-positive early endosomes was followed in SH-SY5Y cells (Figure 1G). The microscopic images reveal that a considerable quantity of oAβ_1-40_-TMR remains at the cell surface compared to oAβ_1-42_-TMR. This effect is more pronounced at 15 min than at 30 min. When endocytosis is inhibited with IPA-3, more oAβ_1-40_-TMR and oAβ_1-42_-TMR remain on the cell surface. Colocalization analysis shows that oAβ_1-42_-TMR colocalizes significantly higher in EEA1-positive early endosomes than oAβ_1-40_-TMR. IPA-3 treatment reduces colocalization for both oAβ_1-40_-TMR and oAβ_1-42_-TMR in the early endosomes (Figure 1H). Internalization follows EEA1-positive early endosomes to Lamp1 vesicles. WB data show a noticeably increased expression of EEA1 with oAβ_1-42_ internalization, and IPA-3-mediated inhibition of endocytosis/internalization reduces EEA1 expression (Figure 1I and J).

Our previous study showed that oAβ_1-42_ internalization follows activated pPAK1-mediated actin-dependent endocytosis (4). The internalization of aggregation-prone oAβ peptides through pPAK1-mediated endocytosis ultimately leads to the buildup of lysosomal vesicles (21). Similarly, we observe here that aggregation-prone oAβ_1-42_ causes a greater extent of Lamp1 expression compared to oAβ_1-40_ (Figure 1K, L). IPA-3 inhibits the pPAK1-dependent endocytosis of oAβ peptides into Lamp1-positive endo-lysosomal vesicles. This leads to decreased levels of Lamp1, with a notable reduction seen in cells treated with oAβ1-42 (Figure 1L). The expression of Lamp1 following the internalization of oAβ_1-40_ is negligible, and the decrease in Lamp1 levels with IPA-3 is similarly inconsequential (Figure 1M).

### pPAK1 regulates Rab3a, which is involved in the repairing of oAβ induced damaged PM

Rho family GTPases Cdc42 and Rac1 activate PAKs, inducing phosphorylation of serine/threonine sites in kinases that regulate downstream targets, which influence actin cytoskeleton dynamics and various forms of endocytosis (29). Endocytosis of oAβ_1-42_ follows activated pPAK1-mediated actin-dependent CIE (4). In this study, we noted an increase in pPAK1 following the internalization of oAβ peptides through WB analysis, further demonstrating pPAK1’s role in oAβ endocytosis (Figure 2A-C). Upregulation of pPAK1 with oAβ_1-42_ is comparatively higher than oAβ_1-40_ (Figure 2C). WB data were quantitatively analyzed to compare pPAK1 levels in the oAβ_1-42_ and oAβ_1-40_ treated neuronal cells, normalizing with the control expression levels (Figure 2C). The results show that IPA-3 inhibits pPAK1 in oAβ_1-42_ treated cells significantly more than in oAβ_1-40_ treated cells (Figure 2C).

**Figure 2:**
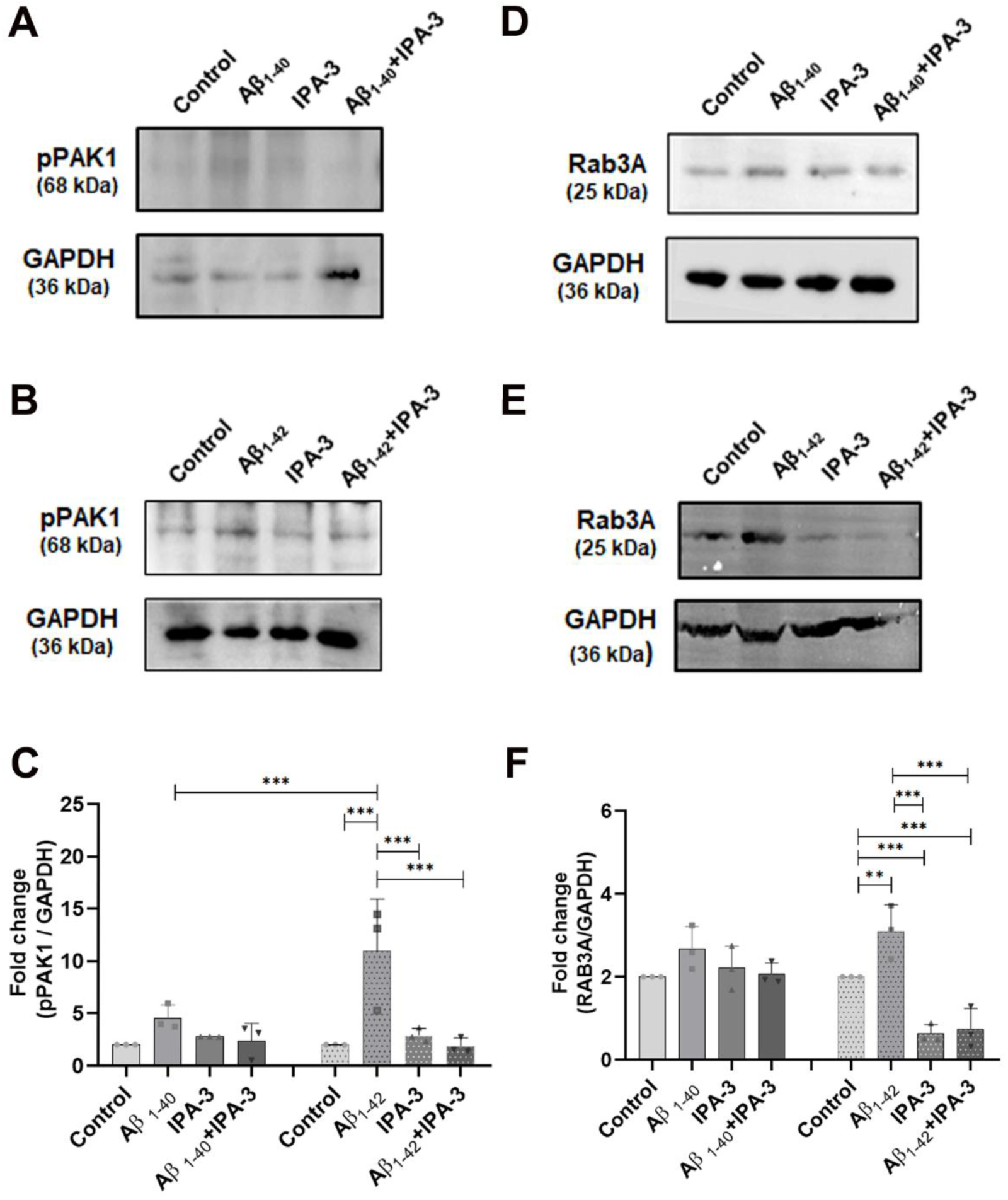
pPAK1 regulates Rab3a expression in the oAβ treated SH-SY5Y cells. WBs of **A, B)** pPAK1 for Aβ _1-40_ and Aβ _1-42_ treated cells with and without IPA3 treatment. Quantification of **C)** pPAK1 levels. WBs of **D, E)** Rab3a for Aβ _1-40_ and Aβ _1-42_ treated cells with and without IPA3 treatment. Quantification of **F)** Rab3a levels. The data are presented as mean ± SD, with *** denoting significance at p ≤ 0.001. The statistical analysis was performed using one-way ANOVA.

In neurons, Cdc42, Rac1, and PAK1 regulate synaptic vesicle trafficking and exocytosis by modulating Rab3 and other Rab family members that facilitate vesicle fusion with the presynaptic membrane (30). Rab3a is a small GTPase protein recognized for its role in PM repair through vesicle fusion (18, 31). Therefore, we investigated whether pPAK1 is associated with Rab3a-mediated PM repair. We observed that pPAK1 helps to regulate the Rab3a expression in response to oAβ exposures to neuronal cells SH-SY5Y and SK-N-SH cells. WB experiments reveal upregulation of Rab3a due to internalization of oAβ peptides (Aβ_1-42_ and Aβ_1-40_) via pPAK1-mediated actin-dependent endocytosis in neuronal cells (Figure 2D-E). Inhibition of pPAK1 by IPA-3 inhibits the expression of Rab3a. Rab3a expressions were quantified and compared between oAβ_1-42_ and oAβ_1-40,_ normalizing the control expression levels (Figure 2F). The result shows that Rab3a expression is significantly higher in oAβ_1-42_ treated cells compared to oAβ_1-40_ (Figure 2F). Studies showed Rab3a-mediated exocytosis plays a role in PM repair (16). The effect of coupled endocytosis and exocytosis is highly relevant in cases of toxic oAβ_1-42_ induced PM damage incidents (4). The effect is visible to a lesser extent in the case of less toxic oligomers of oAβ_1-40_.

### Dynamics of the Rab3a-positive vesicles during PM **repair** induced by oAβ peptides

Exocytosis of lysosomal vesicles plays a major role in the resealing of damaged PM (32). However, the molecular machinery involved in PM repair via lysosomal exocytosis is poorly understood. Localization of the Rab3A was observed on the surface of synaptic vesicles of various types of neurons, including motor, sensory, adrenergic, and cholinergic neurons (33). Translocation of Rab3a-positive vesicles to the PM and their positioning (34, 35) via lysosomal exocytosis plays a significant role during PM repair. Therefore, we have studied the distribution of the Rab3a-positive vesicles at the PM of SH-SY5Y cells in the presence of oAβ peptides (Figure 3). Upon treatment with oAβ peptides, we observed increased localization of Rab3a puncta at the PM in the TIRF field. The density of the Rab3a puncta was higher in oAβ_1-42_ treated cells when compared to oAβ_1-40_ treated and control cells (Figure 3A-D; Movie S1-3). This increase in density suggests fast recycling of Rab3a at the PM (Movie S1-3). To study the dynamics of Rab3a puncta, we analyzed the movement of individual puncta near the PM, with and without oAβ treatments. In oAβ_1-42_ treated cells, Rab3a puncta resided for a significantly shorter duration at the PM (34.2s) when compared to oAβ_1-40_ treated (76.4s) and control cells (71.1s) (Figure 3E-F). The data is plotted as the residence of a single Rab3a puncta in the TIRF field. The change in the fluorescence of a single Rab3a puncta (Figure 3G-I) and the average fluorescence change of Rab3a puncta over time is plotted (Figure 3J-K). In the whole population, fluorescent Rab3a puncta were tracked over time (Figure 3L-N), and the results show an increase in the total and average distance traveled by the Rab3a puncta in oAβ_1-42_ treated cells when compared to oAβ_1-40_ treated and control cells (Figure 3O-P). oAβ_1-42_ treatment increases the overall movement of Rab3a puncta, with shorter residence at the PM for faster recycling of the Rab3a puncta for Rab3a-dependent exocytosis in PM repair.

**Figure 3:**
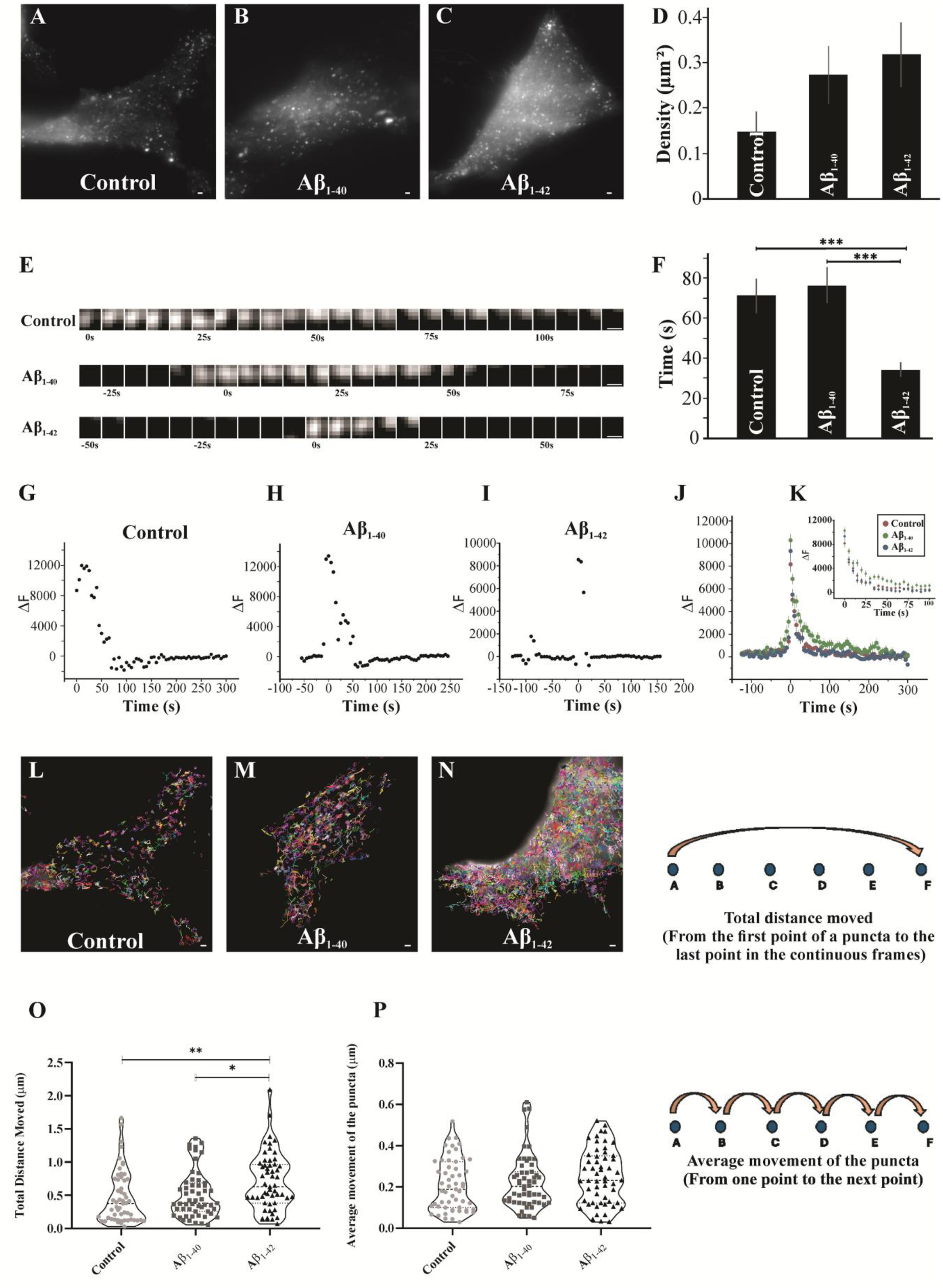
Dynamics of RAB3a positive vesicles near the PM of SH-SY5Y cells. Image of a cell showing EGFP-RAB3a puncta near the PM **(A)** Control, without Aβ treatment, **(B)** with Aβ_1-40_, and **(C)** with Aβ_1-42_ treatment. Scale bar 1 μm. **D)** Quantification of density of EGFP-RAB3a puncta near the PM. Data are expressed as mean ± SEM for at least n=116 puncta (from n=5 cells) for each condition from at least 2 independent experiments. **E)** Time series strip showing dynamics of RAB3a puncta near the PM. Scale bar 0.25μm. **F)** Residence time of RAB3a puncta near the PM with or without Aβ treatment. Graphical representation of fluorescence from single EGFP-RAB3a puncta event as shown in E**, G)** without Aβ treatment, **H)** with Aβ_1-40_, and **I)** with Aβ_1-42_ treatment. **J)** Average of fluorescence from events such as in E showing the dynamics without or with Aβ treatment. **K)** Same as J shown for a period of 100s. Data are expressed as mean ± SEM for n=75 puncta from 5 cells in each case, at least from 2 independent experiments. T-test with *** indicating p ≤ 0.001. **L-N)** Image of a cell showing tracks of EGFP-RAB3a puncta near the PM. **L)** without Aβ treatment, **M)** with Aβ_1-40,_ and **N)** with Aβ_1-42_ treatment. Scale bar 1μm. **O)** Total distance and **P)** average distance travelled by EGFP-RAB3a puncta near the PM as shown in L-N. Data are expressed as mean ± SEM for n=75 puncta from 5 cells in each case, at least from 2 independent experiments.

Additionally, we found that IPA-3 treatment decreases the localization of Rab3a puncta at the PM in oAβ_1-42_ treated cells within the TIRF field (Figure S3A-D). The density of the Rab3a puncta decreases in the presence of IPA-3 in oAβ_1-42_ treated cells (Figure S3E). The dynamics of Rab3a puncta were examined based on the movement of individual puncta close to the PM following IPA-3 mediated inhibition of endocytosis in the oAβ_1-42_ treated cells (Figure S3F). The results indicate that RAB3a puncta reside longer near the PM following IPA-3 treatment (Figure S3G), represented by the residence of a single Rab3a puncta in the TIRF field. Rab3a puncta were tracked over time during IPA-3 treatment (Figure S3H), and the results show a decrease in both the total and average distance traveled by the Rab3a puncta upon the inhibition of oAβ_1-42_ endocytosis by IPA-3 (Figure S3I-J). The results indicate that IPA-3 reduces Rab3a puncta recycling at the PM, inhibiting Rab3a-dependent exocytosis in PM repair.

### Distribution of Rab3a upon oAβ peptides induced PM damage in neuronal cells

Further, we have looked for the distribution of Rab3-dependent lysosomal (Lamp1-positive) vesicles in oAβ peptides treated SH-SY5Y cells. We observed the distribution of significantly higher numbers of Lamp1 positive vesicles colocalized with Rab3a in the periphery of the oAβ_1-42_ treated cells, whereas in the control cells and oAβ_1-40_ treated cells Rab3a positive Lamp1 vesicles are peri-nuclear (Figure 4A-C). IPA-3 treatment results in a significant reduction of Rab3a expression and inhibition of vesicle distribution at the cell periphery in oAβ_1-42_ treated cells (Figure 4C).

**Figure 4:**
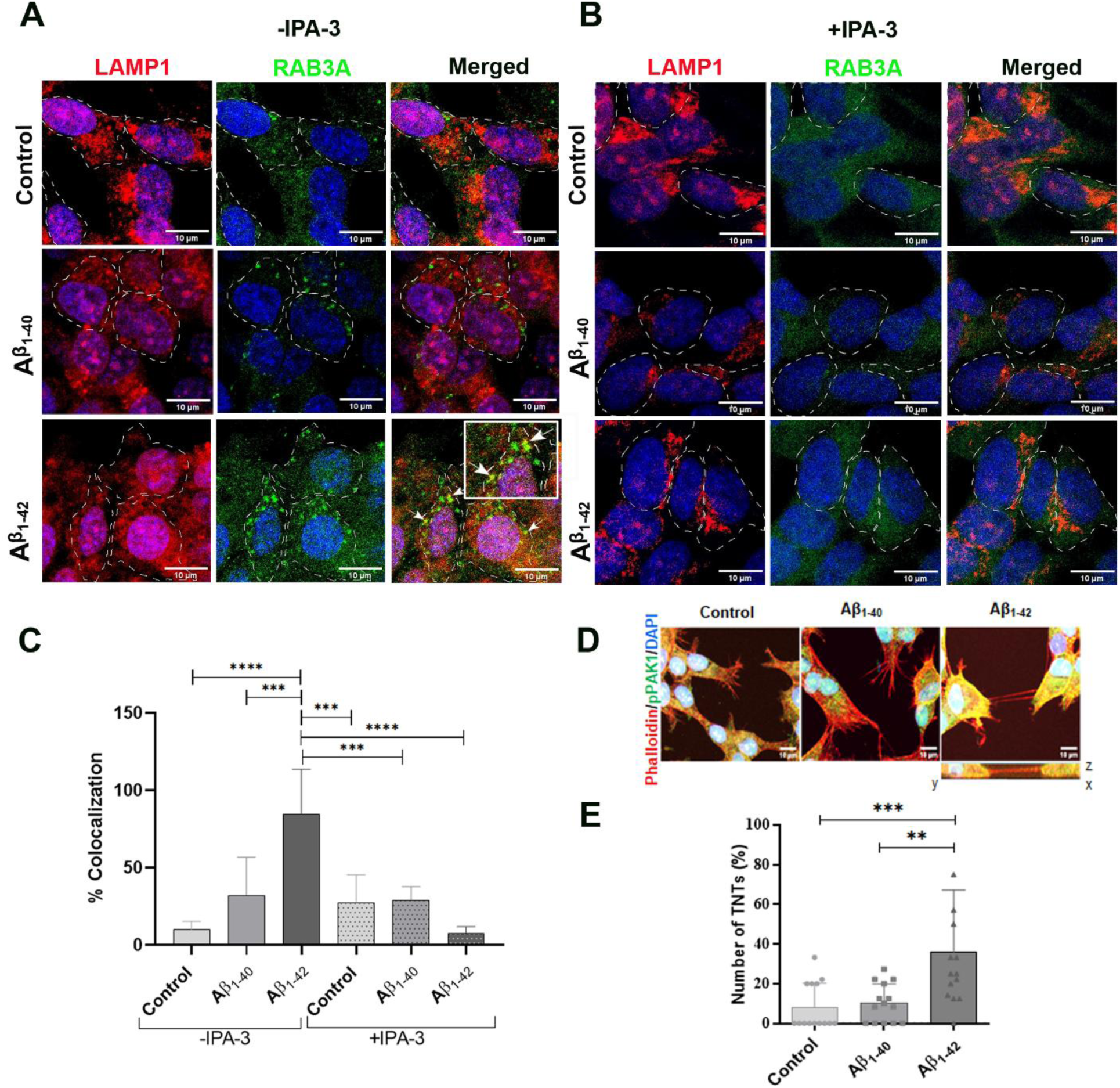
Distribution of Rab3a vesicles in oAβ treated SH-SY5Y cells. **A,B)** Lamp1(red) and Rab3a (green) distributions in control, oAβ_1-40_ and oAβ_1-42_ after 2 hours of treatments, in the absence **(A)** and presence **(B)** of IPA3 pretreated (30 minutes) cells. The distribution of colocalized vesicles is indicated by white arrows in oAβ_1-42_ treated cells, and magnified images are shown in the white box. **C)** Quantification of the percentage of colocalization of Lamp1 positive vesicles with Rab3. Data of average colocalization from 3 independent experiments was presented as mean ± SD. Average **D)** Visualizing TNTs in cells under different conditions: control, with oAβ_1-40_, and oAβ _1-42_ treatment for 2 hours. The lower panel depicts the 3D reconstruction of the TNT image, illustrating it as a suspended or hovering structure positioned between two neighboring cells. This distinctive feature aids in distinguishing TNTs from other neurites. **E)** Quantifying the percentage of TNTs and presenting the data as mean ± SD. The analysis was conducted with the number of images of n = 12, from at least 2 independent experiments. Significance was assessed using one-way ANOVA, and the results indicate *** p ≤ 0.001.

In a previous study (4), our results suggested that membrane nanotubes form due to PM repair in response to oAβ_1-42_ induced PM damage, probably to maintain homeostasis of membrane-actin stress by rapid recycling of vesicles during PM repair dynamics. Damage caused by oAβ_1-42_ induces a repair mechanism via pPAK1 and RAb3a coupled endocytosis-exocytosis at a greater extent with faster dynamics than oAβ_1-40_. We have observed that in the cells treated with oAβ_1-42_, TNT-like structures are higher in numbers compared to oAβ_1-40_, correlating with faster PM repair dynamics (Figure 4D-E). The result reveals that preceding the formation of TNT-like membrane protrusions, PM repair occurs in Aβ-induced PM-damaged neuronal cells.

### Ca^2+^-dependent lysosomal vesicle docking is crucial for PM **repair** and preventing neurotoxicity

Rab3a plays a crucial role in the Ca^2+^-dependent lysosomal exocytosis by facilitating vesicle docking (35, 36). The intensity of Propidium iodide (PI) leakage in the absence of Ca^2+^ upon membrane damage indicates the extent of PM damage (23). Flow cytometry experiments with PI leakage have already been shown in our previous study (4). Here, we performed quantification of PI leakage using a microscopy experiment. We quantified the PI nuclear staining colocalized with Hoechst in the live SH-SY5Y cells treated with oAβ_1-40_ and oAβ_1-42_ (Figure 5A and B). We compared oAβ_1-40_ and oAβ_1-42_ treated cells with the control in two different conditions, such as in the presence (Figure 5A) and absence of Ca^2+^ (Figure 5B), using EGTA buffer as a Ca^2+^ chelator. PI-positive cells were comparatively less in control and oAβ_1-40_ treated cells in both the presence and absence of Ca^2+^. The numbers of PI-positive cells are significantly higher in oAβ_1-42_ treated cells compared to control and oAβ_1-40_ treated cells both in presence and absence of Ca^2+^ (Figure 5C). PI-positive cells are also significantly higher in oAβ_1-42_ treated cells in absence of Ca^2+^, while comparing the Ca^2+^ treated oAβ_1-42_ cells (Figure 5C). It reveals the repair of oAβ peptides-induced membrane damage is Ca^2+^-dependent, and the extent of membrane damage by oAβ_1-40_ is significantly lower compared to oAβ_1-42_ (Figure 5C).

**Figure 5:**
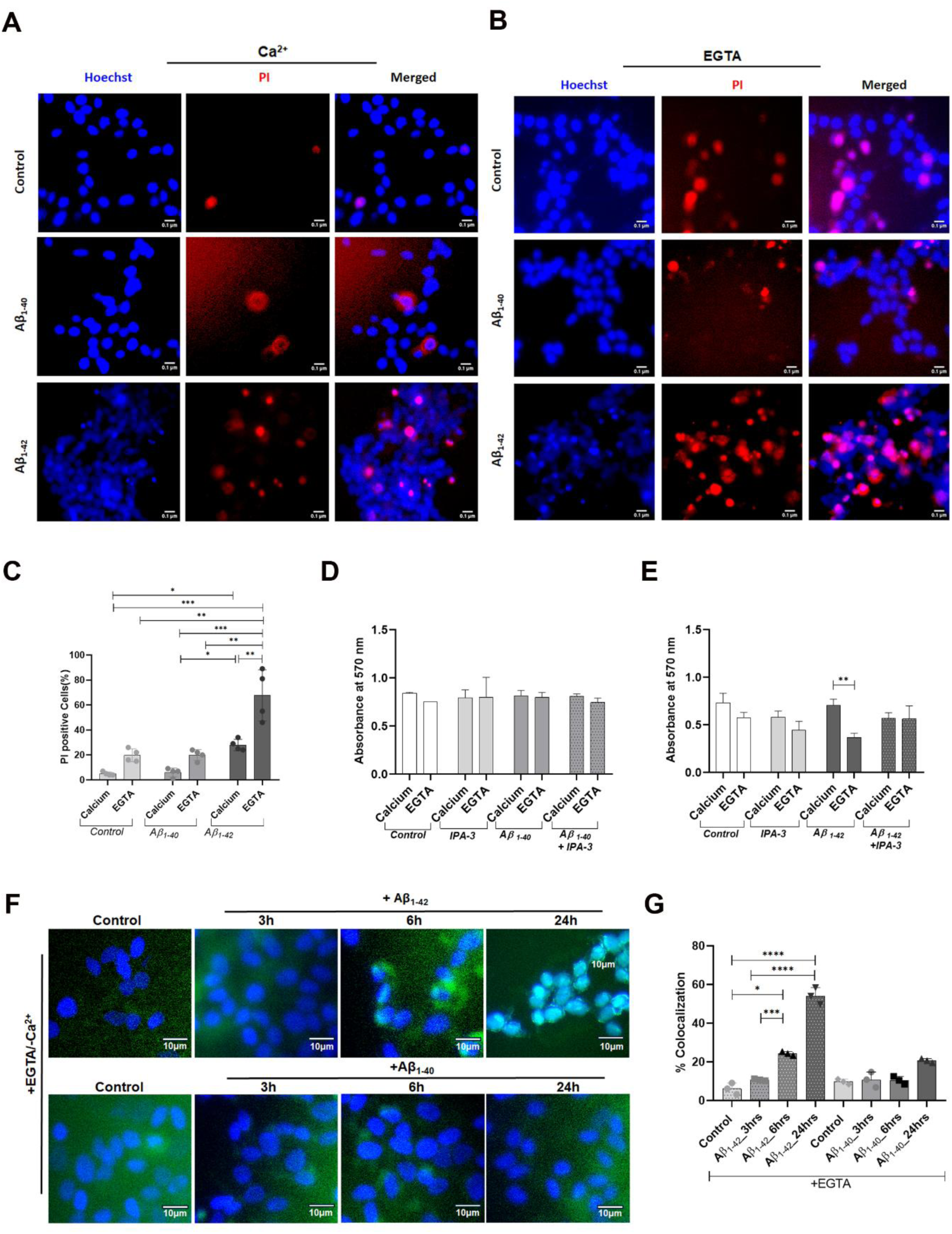
Ca^2+^-dependent PM repair in the oAβ treated SH-SY5Y cells. **A, B)** Propidium iodide (PI) and Hoechst double staining in SH-SY5Y cells after 1-hour treatment with oAβ_1-40_ and oAβ_1-42_ **A)** presence and **B)** absence of Ca^2+^. **C)** Graphical representation of the percentage of PI-positive cells per unit area of cells in both the presence and absence of Ca^2+^ conditions. **D-E)** MTT assay results were obtained **D)** oAβ_1-40_ and **E)** oAβ_1-42_ in presence and absence of IPA-3 for 3 hrs. Data are expressed as mean ± SD, *** p ≤ 0.001. Statistics were analyzed using two-way ANOVA. n=4. **F)** TUNEL assay was performed in oAβ_1-40_ and oAβ_1-42_ treated cells in the presence of EGTA at 3, 6, and 24 hours. **G)** Quantification of percentage of colocalization of DAPI (blue) with anti-BrdU (green) were graphically represented. The image analysis was done from 3 independent experiments. The data is presented as mean ± SD, and significance was observed with **** p ≤ 0.0001. Statistical analysis was conducted using one-way ANOVA.

PM repair plays a crucial role in neuroprotection. MTT assay was done to quantify cell viability in the presence and absence of Ca^2+^ in the oAβ_1-40_ and oAβ_1-42_ treated cells. The relatively non-toxic oligomers of Aβ_1-40_ peptide, which caused insignificant PM damage, show no significant change in cell viability regardless of the presence and absence of Ca^2+^ even after 6 hours of treatment (Figure 5D). Compared to that, oAβ_1-42_ treated cells show a significant reduction in cell viability without Ca^2+^ buffer (Figure 5E). IPA-3, which inhibits PM repair by reducing Rab3a upregulation, attenuates the survival of oAβ_1-42_ treated cells in the presence and absence of Ca^2+^ buffer (Figure 5E). MTT assay is based on mitochondrial activity-regulated metabolic viability. Thus, we have performed a TUNEL assay to monitor genomic DNA break-induced apoptotic cell death. The TUNEL assay reveals a notable rise in green fluorescent labelling (using Alexa Fluor 488 labelled anti-BrdU monoclonal antibody) within the nucleus, attributed to the transfer of BrdUTP to the 3’-hydroxyl ends of DNA double-strand breaks in the oAβ_1-42_ treated cells after 24 hours, with EGTA buffer present (Figure 5F-G). The findings indicate that apoptosis is induced in oAβ_1-42_ treated cells after 24 hours when using EGTA buffer. The TUNEL assay did not show significant DNA double-strand breaks in cells treated with less aggregation-prone oligomers of oAβ_1-40_ (Figure 5F-G). The inhibition of endocytosis by IPA-3, alongside the chelation of Ca²⁺ with EGTA, which prevents exocytic vesicle fusion, leads to apoptosis and significant fluorescence staining in the nuclei of oAβ_1-42_ treated cells, observable as early as 3 hours (Figure S4A-B). Proper exocytic vesicle fusion with Ca²⁺ repairs PM, preventing apoptotic cells in oAβ_1-42_ treated cells after 24 hours (Figure S4C-D). The result reveals a crucial role of PM repair in neuronal cell survival.

### oAβ peptides induced PM damage and its repair in the Rab3a knockdown neuronal cells

Our aim is to inhibit Rab3a expression to understand the role of Rab3a in lysosomal exocytosis-mediated PM repair. Knockdown of Rab3a expression by shRNA was performed in the SH-SY5Y neuronal cells as per the mentioned protocol. Knockdown efficiency was assessed from WBs bands of Rab3a protein from shRab3A transfected cells, compared to pLKO.1 lentiviral vector-transfected cells as a control. The results show a reduction to 40.5 ± 24.4 % Rab3a expression in the knockdown cells up to 5 consecutive passages under puromycin selection (Figure 6A). WB results further revealed that Rab3a knockdown caused a reduction in pPAK1 levels (Figure 6B). In the Rab3a knockdown cells, PM damage was assessed by determining PI leakage into the cells in the presence of Ca^2+^ buffer using fluorescence microscopy images. We quantified the PI nuclear staining colocalized with Hoechst in the pLKO.1 and shRab3a transfected live SH-SY5Y cells treated with oAβ_1-42_ for 30 minutes and 1 hour (Figure 6C and D). The quantification of the microscopic images shows that the percentage of PI-positive cells is significantly higher in the oAβ_1-42_ treated shRab3a knockdown cells, compared to the pLKO.1 transfected cell in the presence of Ca^2+^ buffer (Figure 6E). The results suggest the essential role of Rab3a in the repair of oAβ_1-42_ induced PM damage. Further, we have observed that Rab3a-dependent PM repair is essential for the viability of neuronal cells. Cell viability assay by MTT has revealed that Rab3a knockdown cells are susceptible to toxic effects of oAβ_1-42,_ and significant cell death was observed within 6 hours of treatment. Where pLKO.1 transfected cell shows no significant cell viability upon oAβ_1-42_ treatment for 6 hours (Figure 6F). The results suggest a significant role of Rab3a in PM repair and the prevention of oAβ_1-42_ toxicity-induced neuronal cell death.

**Figure 6:**
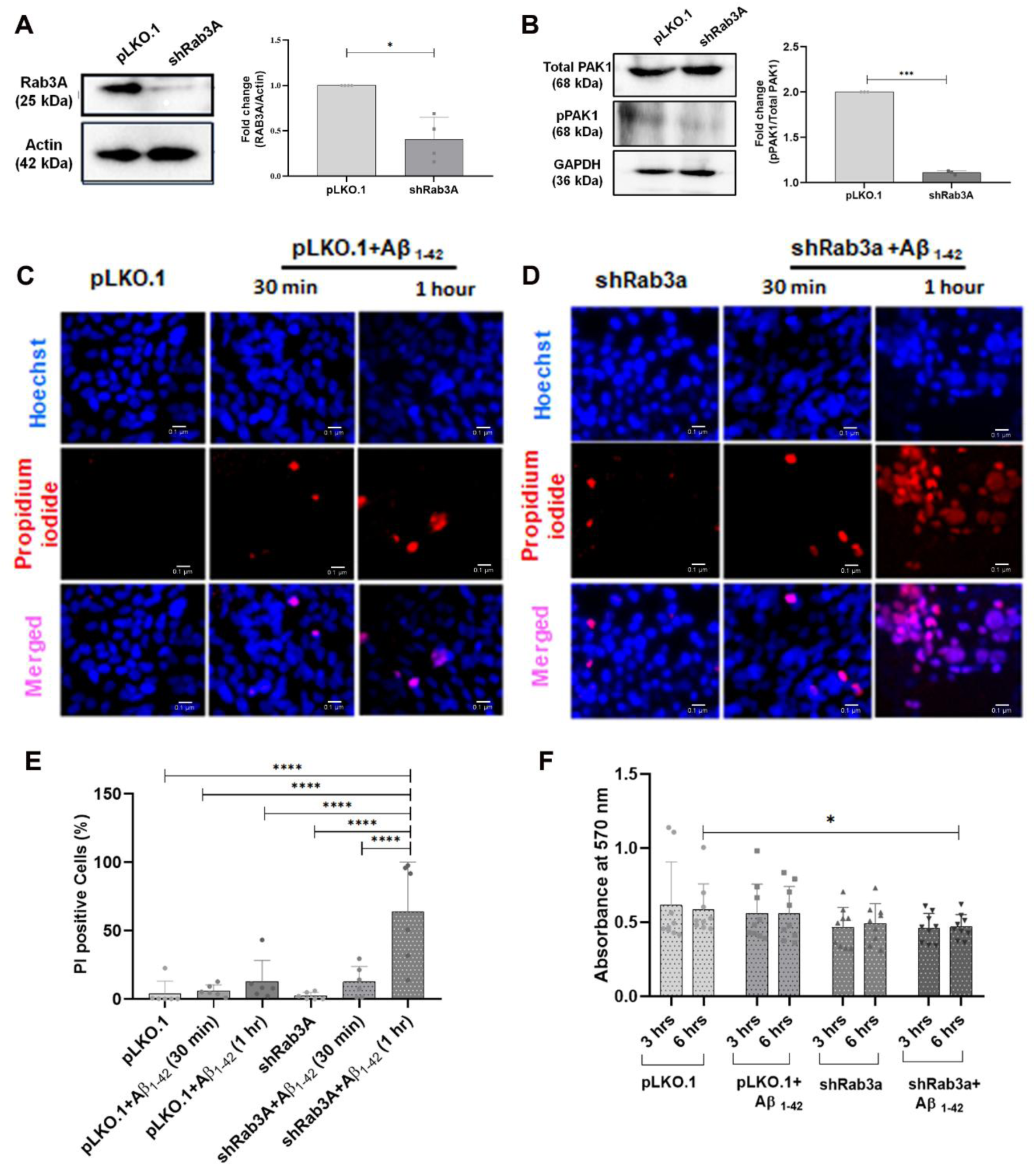
PM repair in the Rab3a knockdown SH-SY5Y cells. **A)** Knockdown efficiency was assessed through the WB experiment conducted on pLKO.1 lentiviral vector-transfected cells as a control and cells with Rab3a knockdown (shRab3A). **B)** WB of total-PAK1 and pPAK1 on pLKO.1 lentiviral vector-transfected cells as a control and cells with Rab3a knockdown (shRab3A). The quantifications are presented as mean ± SD, with *** indicating p ≤ 0.001. Statistical analysis employed a t-test, and n=3. **C-D)** Dual nuclear staining using Propidium iodide and Hoechst was performed on **C)** pLKO.1 and **D)** shRab3a transfected cells, treated with Aβ_1-42_ for 30 minutes and 1 hour. **E)** Graphical representation of PI-positive cells per unit area percentage in both pLKO.1 and shRab3a transfected cells. **F)** Cell viability by MTT assay was performed in pLKO.1 and shRab3a transfected cells, treated with and without Aβ_1-42_ for 3 and 6 hours. The data is expressed as mean ± SD, with *** indicating p ≤ 0.001. One-way ANOVA was used for statistical analysis. Statistical analysis involved ordinary two-way ANOVA, and the sample size was 9 from three independent experiments (n=3).

## Discussion

Mechanisms that induce AD pathogenesis are largely unknown. Toxic aggregates of Aβ peptides are considered critical initiator that triggers the progress of pathogenesis (37, 38). Aβ-peptides generate extracellularly due to cleavage of transmembrane protein APP (amyloid precursor protein) at PM via β- and γ-secretase enzymes (39). Interaction of these peptides with PM is considered as the main site for initiating the formation of toxic aggregates (7). The membrane environment triggers the transition of native monomeric Aβ to an unfolded conformation, which acts as nucleation to enhance self-aggregation, and eventually to fibrillar aggregates (40). Recent studies indicated soluble oligomers of Aβ-peptides are more toxic than higher-ordered fibrils and plaques (8, 41). Several studies have reported in various model systems that oAβ caused membrane disruptions and lipid distortions. Changes in membrane toxicities caused by toxic Aβ-peptide oligomers trigger neuronal toxicities, leading to the gradual progression of pathology and, eventually, neuronal death (42). Mature neurons strive to survive and employ various strategies to prevent cell death. When toxic oligomers damage the PM, neurons initiate a repairing process to fix the damage caused to the membrane (4, 5). However, it is not widely understood how damaged membranes induced by oAβ initiate PM repair cascades (43). Continuous damage caused by oAβ peptides can lead to neuronal death due to neurotoxicity, as it results in defects in PM repair (44). Therefore, it is important to understand the exact mechanism of PM repair cascades initiated by oAβ peptides.

This study first shows that oligomers of Aβ peptides trigger a PM repair response by promoting the fusion of Rab3a-positive vesicles at the PM, linked to pPAK1-dependent CIE. Our results demonstrated that aggregation-prone Aβ_1-42_ peptide, which forms higher orders of aggregates compared to Aβ_1-40_, caused a significant extent of PM damage-induced neurotoxicity (Figure 7). Mutations that generate aggregation-prone Aβ peptides induce toxicities related to lipid oxidation, membrane pores formation, and PM damage (3, 40). It is known in the literature that higher molecular weight oAβ_1-42_ induces larger PM damage and neurotoxicity (40). One of the studies has shown that similar to membrane pore-forming bacterial toxin streptolysin-O, oAβ_1-42_ induced PM damage induces PM repair response (5). We have observed that the aggregation-prone oAβ_1-42_, which generates more damage to PM, internalizes to endo-lysosomes to a significantly higher extent than Aβ_1-40_.

**Figure 7:**
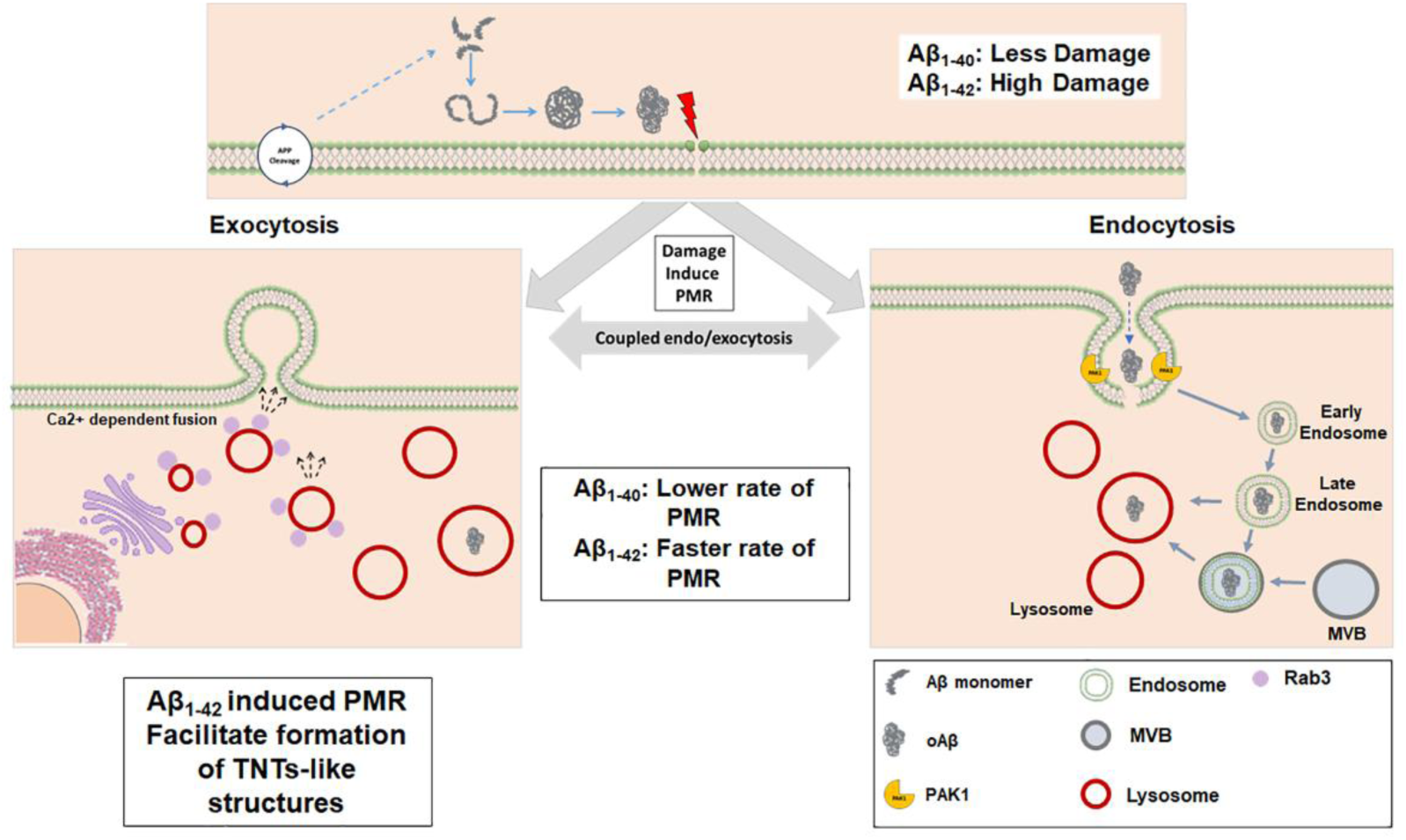
The extracellular oAβ induces PM damage: oAβ_1-40_ causes less damage, while oAβ_1-42_ causes higher damage. The PM repair occurs via the Rab3-dependent recycling of Lamp1-positive lysosomal vesicles coupled with pPAK1-mediated massive endocytosis of the oAβ peptides. PM repair, in response to oAβ1-42 induced damage, occurs at a much faster rate. pPAK1 regulates actin-membrane stress via controlling CIE and by forming long-stretched fibrillary actin conduits or TNT-like structures during rapid PM repair.

Internalization of toxic aggregates of oAβ with damaged membranes alters the endo-lysosomal pathway (21). Toxicities in endo-lysosomal pathways are one of the mechanisms of gradual pathology development in AD. We have shown that oAβ_1-42_ induced PM damage initiates massive endocytosis through an actin-dependent pPAK1-mediated CIE pathway (4). On-demand, rapid PM repair in response to oAβ_1-42_ treatment may destabilize membrane-actin tension. CIE plays a crucial role in maintaining membrane tension and cellular homeostasis (9). A previous study reported that Aβ-aggregates follow actin-dependent CIE through Rho GTPase regulated pathway, which helps to maintain PM stress (10). Moreover, the involvement of pPAK1-dependent actin-cytoskeleton modulation during rapid Rab3a positive lysosomal vesicle turnover facilitates the modulation of membrane-actin protrusions. As a result, formation of TNT-like cell-to-cell conduits. In a previous study, we reported pPAK1-dependent biogenesis of TNT-like structures upon oAβ_1-42_-dependent PM repair (4). This study reveals that rapid PM repair occurs through Rab3a vesicle fusion when treated with oAβ_1-42_ peptides, which correlates with developing TNT-like structures. In comparison, oAβ_1-40_ leads to reduced PM damage and slower repair rates, resulting in fewer TNT-like structures.

Increased levels of Rab3a play a significant role in positioning lysosomes and repairing damaged PM. In our single-particle tracking experiments, we observed clearly faster recruitment of EGFP-Rab3a positive vesicles at the PM upon oAβ_1-42_ treatment than oAβ_1-40_. oAβ_1-42_ treatment induced faster Rab3a dynamics and the increase in density suggests faster recycling of Rab3a puncta for Rab3a-dependent exocytosis in PM repair. PM damage triggers repair response by Ca^2+^-dependent exocytosis of lysosomes followed by damaged membrane endocytosis (24). Our previous study showed that oAβ_1-42_ induced PM damage triggers PM repair by lysosomal exocytosis coupled with massive endocytosis (4). This study revealed that pPAK1-dependent endocytosis coupled with Rab3a-dependent lysosomal exocytosis. Inhibition of pPAK1 by specific inhibitor IPA-3 treatment results in a significant reduction of Rab3a expression and inhibition of vesicle distribution at the cell periphery in oAβ_1-42_ treated cells. Conversely, the study showed that inhibiting Rab3a expression reduces pPAK1 levels. The study does not elucidate the molecular players that could influence pPAK1-mediated regulation of Rab3a expression and the positioning of Rab3-vesicle at the PM during exocytosis (30). Research indicates that Cdc42, Rac1, and activated PAKs in neuronal cells can influence the fusion of Rab3-positive presynaptic exocytic vesicles. The Rho family GTPases, namely Cdc42 and Rac1, activate PAKs that phosphorylate serine and threonine residues in kinases, subsequently impacting actin dynamics, endocytosis, as well as the trafficking and exocytosis of synaptic vesicles (29).

Overall, Aβ production is influenced by APP trafficking and its concomitant localization with affiliated secretases in endosomes. Rab proteins, along with other trafficking factors, are involved in this process. The dysregulation of these trafficking factors may impact the development of AD pathogenesis by contributing to the production of Aβ (45). *RAB23, RAB8a/RAB8, and RAB3a*, are highly expressed in the brain and engage in neural development, synaptic vesicle exocytosis, and neurite growth (46, 47). Dysregulation of AD-related Rab GTPases collapse endosomal, lysosomal, recycling, and autophagic pathways. Rab3a mainly affects APP assembly into recycling vesicles during anterograde transport (48), which affects APP cleavage and Aβ generation (49). In the context of AD pathogenesis in relation to the PM repair, a study from 2016 (19) showed that silencing of Rab3a affects lysosome positioning, terminating the destruction of lysosomes, thereby inhibiting PM repair. They identified that this is mediated by the effector Slp4-a, and NMHC II. A complex of Rab3a-Slp4-a-NMHCIIA is involved in the positioning of lysosomes at the periphery and lysosome exocytosis.

Several studies have reported the role of Rab3a in AD. Rab3a has a pivotal role in Ca^2+^-dependent exocytosis and vesicle docking of neurotransmitters in synapses (36, 50). Vesicles containing higher levels of Rab3a recycle more actively than vesicles that are stationary (51). Disruption of anterograde transport increases Rab3 levels, correlated with age-related cognitive decline (52). Among four isoforms of Rab3 (Rab3a, Rab3b, Rab3c, and Rab3d), Rab3a has a pivotal role in the Ca^2+^-dependent exocytosis and vesicle docking in synapses (36). Lysosomal network protein screening found increased Rab3 and Rab7 levels in AD patient CSF (53). These observations indicate Rab3a may have a role in the early stages of development of the AD.

The interaction of toxic oligomers of Aβ peptides with the PM triggers membrane alterations/damage that could initiate progressive neurotoxicity in AD. However, neurons strive to survive and employ various strategies to prevent cell death. When toxic oligomers damage the PM, neurons initiate a repairing process to fix the damage caused to the membrane. However, only a few studies have been done on PM repair in response to Aβ aggregates-induced PM damage (4, 5, 17).

This study illustrates how pPAK-mediated endocytosis affects Rab3a-mediated vesicle fusion at the PM in response to damage caused by Aβ aggregates. We show for the first time the process of pPAK1-dependent endocytosis rate and Rab3a-dependent vesicle fusion at the PM using a TIRF microscope in response to two different types of peptides, the aggregate-prone oAβ_1-42_ and the normal physiological oAβ_1-40_. Compared to the less toxic oAβ_1-40_, the aggregation-prone oAβ_1-42_ significantly induces PM repair cascades. The rapid response of PM repair triggered by PM damage from aggregation-prone oAβ_1-42_ enhances actin modulation and fosters the formation of TNT-like connections between distant cells, likely aiding in the maintenance of membrane stability during homeostasis. pPAK1 is known to manage actin-membrane stress by regulating CIE. These TNT-like connections may enable cell-to-cell propagation (9). IPA-3, a non-ATP competitive inhibitor targeting pPAK1, disrupts the internalization of oAβ, hinders Rab3a-mediated PM repair, and prevents the development of TNT-like actin-membrane protrusions. Inhibition in Rab3a expression using shRNA causes defects in PM repair, which leads to neurotoxicity. The study showed the relevance of Rab3a pathway in the context of oAβ_1-42_ induced recycling of vesicles and endolysosomal toxicities in association with PM repair in the context of AD. The study will open up a new understanding of AD pathogenesis, how interactions of Aβ aggregates at the membrane level, and its relation to PM damage involved in the intercellular spread of pathogenic aggregates. A novel therapeutics route could be developed targeting the molecules related to PM repair pathways in AD.

## Author Credit Statement

SN and DKV conceptualized the work; DKV and PV performed the TIRF-microscopy experiments. NRG and SN designed the work; DKV, SPS, and PH performed the WBs and other experiments; DKV and JMG performed shRNA knockdown experiments. DKV, PV, NRG, and SN validated and investigated all the formal analyses; NRG, AK, and SN provided methodology support, visualization, and interpretation of data; DKV, PV, NRG, and SN wrote the paper, taking valuable inputs from all the authors.

## ACKNOWLEDGEMENTS

DKV thanks the Manipal Academy of Higher Education for the TMA Pai fellowship. SN thanks the Science and Engineering Research Board of India for the SERB-SRG (#SRG/2021/001315); the Indian Council of Medical Research of India (#5/4-5/Ad-hoc/Neuro/216/2020-NCD-I and #IIRP-2023-0084), DST FIST grant #SR/FST/LS-I/2018/121, and the Intramural fund of Manipal Academy of Higher Education, Manipal, India for research funding. NRG’s lab was supported by Indian Institute of Science start-up grants, Indian Council of Medical Research (ICMR) – Grant-in-aid scheme, Department of Biotechnology(DBT) - Ramalingaswami grant, SERB-SRG grant, Longevity India Initiative, Novo Nordisk Foundation and Infosys young investigator grant. PV is supported by the Prime Minister’s Research Fellowship (PMRF). We thank Ms. B. Suma for the confocal microscopy, the JNCASR confocal facility, and the Bioimaging facility of the Indian Institute of Science, Bangalore.

## References

1. Guo, T., Zhang, D., Zeng, Y., Huang, T. Y., Xu, H., and Zhao, Y. (2020) Molecular and cellular mechanisms underlying the pathogenesis of Alzheimer’s disease. Mol. Neurodegener. 15, 40

2. Hampel, H., Hardy, J., Blennow, K., Chen, C., Perry, G., Kim, S. H., Villemagne, V. L., Aisen, P., Vendruscolo, M., Iwatsubo, T., Masters, C. L., Cho, M., Lannfelt, L., Cummings, J. L., and Vergallo, A. (2021) The Amyloid-β Pathway in Alzheimer’s Disease. Mol. Psychiatry. 26, 5481–5503

3. Sengupta, U., Nilson, A. N., and Kayed, R. (2016) The Role of Amyloid-β Oligomers in Toxicity, Propagation, and Immunotherapy. EBioMedicine. 6, 42–49

4. Dilna, A., Deepak, K. V., Damodaran, N., Kielkopf, C. S., Kagedal, K., Ollinger, K., and Nath, S. (2021) Amyloid-β induced membrane damage instigates tunneling nanotube-like conduits by p21-activated kinase dependent actin remodulation. Biochim. Biophys. Acta - Mol. Basis Dis. 10.1016/j.bbadis.2021.166246

5. Julien, C., Tomberlin, C., Roberts, C. M., Akram, A., Stein, G. H., Silverman, M. A., and Link, C. D. (2018) In vivo induction of membrane damage by β-amyloid peptide oligomers. Acta Neuropathol. Commun. 6, 131

6. Rangachari, V., Dean, D. N., Rana, P., Vaidya, A., and Ghosh, P. (2018) Cause and consequence of Aβ - Lipid interactions in Alzheimer disease pathogenesis. Biochim. Biophys. acta. Biomembr. 1860, 1652–1662

7. Valappil, D. K., Mini, N. J., Dilna, A., and Nath, S. (2022) Membrane interaction to intercellular spread of pathology in Alzheimer’s disease. Front. Neurosci. 16, 936897

8. Gouras, G. K., Tampellini, D., Takahashi, R. H., and Capetillo-Zarate, E. (2010) Intraneuronal β-amyloid accumulation and synapse pathology in Alzheimer’s disease. Acta Neuropathol. 10.1007/s00401-010-0679-9

9. Thottacherry, J. J., Kosmalska, A. J., Kumar, A., Vishen, A. S., Elosegui-Artola, A., Pradhan, S., Sharma, S., Singh, P. P., Guadamillas, M. C., Chaudhary, N., Vishwakarma, R., Trepat, X., Del Pozo, M. A., Parton, R. G., Rao, M., Pullarkat, P., Roca-Cusachs, P., and Mayor, S. (2018) Mechanochemical feedback control of dynamin independent endocytosis modulates membrane tension in adherent cells. Nat. Commun. 9, 4217

10. Wesén, E., Lundmark, R., and Esbjörner, E. K. (2020) Role of Membrane Tension Sensitive Endocytosis and Rho GTPases in the Uptake of the Alzheimer’s Disease Peptide Aβ(1-42). ACS Chem. Neurosci. 11, 1925–1936

11. Mandrekar, S., Jiang, Q., Lee, C. Y. D., Koenigsknecht-Talboo, J., Holtzman, D. M., and Landreth, G. E. (2009) Microglia mediate the clearance of soluble Abeta through fluid phase macropinocytosis. J. Neurosci. Off. J. Soc. Neurosci. 29, 4252–4262

12. Doherty, G. J., and McMahon, H. T. (2009) Mechanisms of endocytosis. Annu. Rev. Biochem. 78, 857–902

13. 13. Jacob, T., Broeke, C. Van den, Waesberghe, C. Van, Troys, L. Van, and Favoreel, H. W. (2015) Pseudorabies virus US3 triggers RhoA phosphorylation to reorganize the actin cytoskeleton. J. Gen. Virol. 96, 2328–2335

14. Raghavan, A., Rao, P., Neuzil, J., Pountney, D. L., and Nath, S. (2021) Oxidative stress and Rho GTPases in the biogenesis of tunnelling nanotubes: implications in disease and therapy. Cell. Mol. Life Sci. 79, 36

15. Abounit, S., Wu, J. W., Duff, K., Victoria, G. S., and Zurzolo, C. (2016) Tunneling nanotubes: A possible highway in the spreading of tau and other prion-like proteins in neurodegenerative diseases. Prion. 10.1080/19336896.2016.1223003

16. Raghavan, A., Kashyap, R., Sreedevi, P., Jos, S., Chatterjee, S., Alex, A., D’Souza, M. N., Giridharan, M., Muddashetty, R., Manjithaya, R., Padavattan, S., and Nath, S. (2024) Astroglia proliferate upon the biogenesis of tunneling nanotubes via α-synuclein dependent transient nuclear translocation of focal adhesion kinase. iScience. 27, 110565

17. Bulgart, H. R., Perez, M. A. L., Tucker, A., Giarrano, G. N., Banford, K., Miller, O., Bonser, S. W. G., Wold, L. E., Scharre, D., and Weisleder, N. (2024) Plasma membrane repair defect in Alzheimer’s disease neurons is driven by the reduced dysferlin expression. FASEB J. 38, e70099

18. Vieira, O. V (2018) Rab3a and Rab10 are regulators of lysosome exocytosis and plasma membrane repair. Small GTPases. 9, 349–351

19. Encarnação, M., Espada, L., Escrevente, C., Mateus, D., Ramalho, J., Michelet, X., Santarino, I., Hsu, V. W., Brenner, M. B., Barral, D. C., and Vieira, O. V (2016) A Rab3a-dependent complex essential for lysosome positioning and plasma membrane repair. J. Cell Biol. 213, 631–640

20. Nath, S., Agholme, L., Kurudenkandy, F. R., Granseth, B., Marcusson, J., and Hallbeck, M. (2012) Spreading of Neurodegenerative Pathology via Neuron-to-Neuron Transmission of -Amyloid. J. Neurosci. 10.1523/JNEUROSCI.0615-12.2012

21. Domert, J., Rao, S. B., Agholme, L., Brorsson, A.-C., Marcusson, J., Hallbeck, M., and Nath, S. (2014) Spreading of amyloid-β peptides via neuritic cell-to-cell transfer is dependent on insufficient cellular clearance. Neurobiol. Dis. 10.1016/j.nbd.2013.12.019

22. Boucher, E., Goldin-Blais, L., Basiren, Q., and Mandato, C. A. (2019) Actin dynamics and myosin contractility during plasma membrane repair and restoration: does one ring really heal them all? Curr. Top. Membr. 84, 17–41

23. Corrotte, M., Castro-Gomes, T., Koushik, A. B., and Andrews, N. W. (2015) Approaches for plasma membrane wounding and assessment of lysosome-mediated repair responses. in Methods in cell biology, pp. 139–158, Elsevier, 126, 139–158

24. Andrews, N. W., and Corrotte, M. (2018) Plasma membrane repair. Curr. Biol. 28, R392– R397

25. 25. Valappil, D. K., Raghavan, A., and Nath, S. (2022) Detection and Quantification of Tunneling Nanotubes Using 3D Volume View Images. JoVE (Journal Vis. Exp.

26. Gandasi, N. R., and Barg, S. (2014) Contact-induced clustering of syntaxin and munc18 docks secretory granules at the exocytosis site. Nat. Commun. 5, 3914

27. Veerabhadraswamy, P., Lata, K., Dey, S., Belekar, P., Kothegala, L., Mangala Prasad, V., and Gandasi, N. R. (2024) Comparison of localization and release of multivesicular bodies and secretory granules in islet cells: Dysregulation during type-2 diabetes. J. Extracell. Biol. 3, e70014

28. Paul, S., Pallavi, A., and Gandasi, N. R. (2024) Exploring the potential of pheophorbide A, a chlorophyll-derived compound in modulating GLUT for maintaining glucose homeostasis. Front. Endocrinol. (Lausanne). 15, 1330058

29. Kumari, S., Mg, S., and Mayor, S. (2010) Endocytosis unplugged: multiple ways to enter the cell. Cell Res. 20, 256–275

30. Nik Akhtar, S., Bunner, W. P., Brennan, E., Lu, Q., and Szatmari, E. M. (2023) Crosstalk between the Rho and Rab family of small GTPases in neurodegenerative disorders. Front. Cell. Neurosci. 17, 1084769

31. Geppert, M., Goda, Y., Stevens, C. F., and Südhof, T. C. (1997) The small GTP-binding protein Rab3A regulates a late step in synaptic vesicle fusion. Nature. 387, 810–814

32. Tancini, B., Buratta, S., Delo, F., Sagini, K., Chiaradia, E., Pellegrino, R. M., Emiliani, C., and Urbanelli, L. (2020) Lysosomal Exocytosis: The Extracellular Role of an Intracellular Organelle. Membranes (Basel). 10.3390/membranes10120406

33. Li, J. Y., Jahn, R., Hou, X. E., Kling-Petersen, A., and Dahlström, A. (1996) Distribution of Rab3a in rat nervous system: comparison with other synaptic vesicle proteins and neuropeptides. Brain Res. 706, 103–112

34. Raiborg, C., and Stenmark, H. (2016) Plasma membrane repairs by small GTPase Rab3a. J. Cell Biol. 213, 613–615

35. Corrotte, M., and Castro-Gomes, T. (2019) Lysosomes and plasma membrane repair, pp. 1–16, 10.1016/bs.ctm.2019.08.001

36. Coleman, W. L., Bill, C. A., and Bykhovskaia, M. (2007) Rab3a deletion reduces vesicle docking and transmitter release at the mouse diaphragm synapse. Neuroscience. 148, 1–6

37. Hallbeck, M., Nath, S., and Marcusson, J. (2013) Neuron-to-neuron transmission of neurodegenerative pathology. Neurosci. a Rev. J. bringing Neurobiol. Neurol. psychiatry. 19, 560–566

38. Morris, G. P., Clark, I. A., and Vissel, B. (2014) Inconsistencies and Controversies Surrounding the Amyloid Hypothesis of Alzheimer’s Disease. Acta Neuropathol. Commun. 2, 135

39. Zhang, Y., Thompson, R., Zhang, H., and Xu, H. (2011) APP processing in Alzheimer’s disease. Mol. Brain. 4, 3

40. Yasumoto, T., Takamura, Y., Tsuji, M., Watanabe-Nakayama, T., Imamura, K., Inoue, H., Nakamura, S., Inoue, T., Kimura, A., Yano, S., Nishijo, H., Kiuchi, Y., Teplow, D. B., and Ono, K. (2019) High molecular weight amyloid β(1-42) oligomers induce neurotoxicity via plasma membrane damage. FASEB J. Off. Publ. Fed. Am. Soc. Exp. Biol. 33, 9220–9234

41. Narayan, P., Ganzinger, K. A., McColl, J., Weimann, L., Meehan, S., Qamar, S., Carver, J. A., Wilson, M. R., St. George-Hyslop, P., and Dobson, C. M. (2013) Single molecule characterization of the interactions between amyloid-β peptides and the membranes of hippocampal cells. J. Am. Chem. Soc. 135, 1491–1498

42. Tolar, M., Hey, J., Power, A., and Abushakra, S. (2021) Neurotoxic Soluble Amyloid Oligomers Drive Alzheimer’s Pathogenesis and Represent a Clinically Validated Target for Slowing Disease Progression. Int. J. Mol. Sci. 10.3390/ijms22126355

43. Drolle, E., Negoda, A., Hammond, K., Pavlov, E., and Leonenko, Z. (2017) Changes in lipid membranes may trigger amyloid toxicity in Alzheimer’s disease. PLoS One. 12, e0182194

44. Bulgart, H., Lopez Perez, M., Tucker, A., Usmani, M., and Weisleder, N. (2023) Neuronal Membrane Repair Defect as a Contributor to Amyloid Beta Pathogenesis in Alzheimer’s Disease. Physiology. 38, 5731327

45. Jiang, S., Li, Y., Zhang, X., Bu, G., Xu, H., and Zhang, Y. (2014) Trafficking regulation of proteins in Alzheimer’s disease. Mol. Neurodegener. 9, 6

46. Corbeel, L., and Freson, K. (2008) Rab proteins and Rab-associated proteins: major actors in the mechanism of protein-trafficking disorders. Eur. J. Pediatr. 167, 723–729

47. Gao, Y., Wilson, G. R., Stephenson, S. E. M., Bozaoglu, K., Farrer, M. J., and Lockhart, P. J. (2018) The emerging role of Rab GTPases in the pathogenesis of Parkinson’s disease. Mov. Disord. 33, 196–207

48. Szodorai, A., Kuan, Y.-H., Hunzelmann, S., Engel, U., Sakane, A., Sasaki, T., Takai, Y., Kirsch, J., Müller, U., Beyreuther, K., Brady, S., Morfini, G., and Kins, S. (2009) APP anterograde transport requires Rab3A GTPase activity for assembly of the transport vesicle. J. Neurosci. Off. J. Soc. Neurosci. 29, 14534–14544

49. Udayar, V., Buggia-Prévot, V., Guerreiro, R. L., Siegel, G., Rambabu, N., Soohoo, A. L., Ponnusamy, M., Siegenthaler, B., Bali, J., Simons, M., Ries, J., Puthenveedu, M. A., Hardy, J., Thinakaran, G., and Rajendran, L. (2013) A paired RNAi and RabGAP overexpression screen identifies Rab11 as a regulator of β-amyloid production. Cell Rep. 5, 1536–1551

50. Masumoto, N., Sasaki, T., Tahara, M., Mammoto, A., Ikebuchi, Y., Tasaka, K., Tokunaga, M., Takai, Y., and Miyake, A. (1996) Involvement of Rabphilin-3A in cortical granule exocytosis in mouse eggs. J. Cell Biol. 135, 1741–1747

51. Star, E. N., Newton, A. J., and Murthy, V. N. (2005) Real-time imaging of Rab3a and Rab5a reveals differential roles in presynaptic function. J. Physiol. 569, 103–117

52. 52. Kimura, N., Okabayashi, S., and Ono, F. (2012) Dynein dysfunction disrupts intracellular vesicle trafficking bidirectionally and perturbs synaptic vesicle docking via endocytic disturbances a potential mechanism underlying age-dependent impairment of cognitive function. Am. J. Pathol. 180, 550–561

53. Armstrong, A., Mattsson, N., Appelqvist, H., Janefjord, C., Sandin, L., Agholme, L., Olsson, B., Svensson, S., Blennow, K., Zetterberg, H., and Kågedal, K. (2014) Lysosomal network proteins as potential novel CSF biomarkers for Alzheimer’s disease. Neuromolecular Med. 16, 150–160

