## Supplementary Text for "p21-activated kinase regulates Rab3a vesicles to repair plasma membrane damage caused by Amyloid-β oligomers"

**Short title:** Plasma membrane repair in response to Amyloid-β oligomers

**Keywords:**

Plasma Membrane Repair, Rab3a, Amyloid-β, p21-activates kinase, Lysosomal exocytosis, Clathrin-independent endocytosis, Tunneling nanotubes,

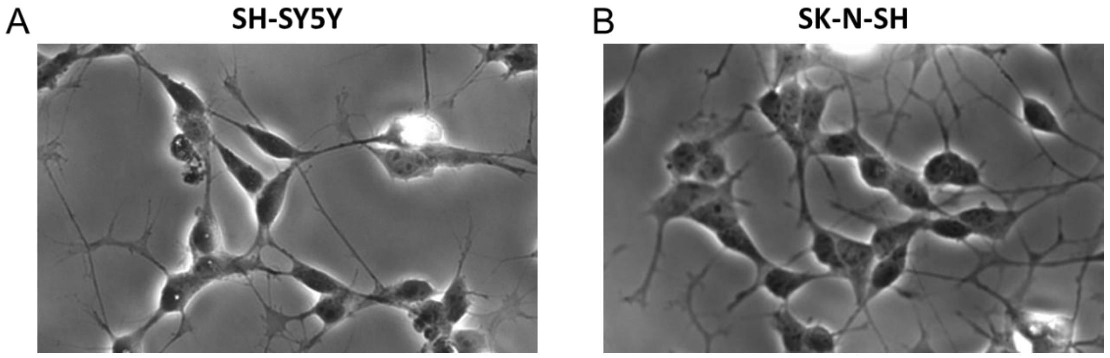


**Figure S 1:** Human neuroblastoma cells were used for the study A) SH-SY5Y cells B) SK-N-SH cells. Both of the cells are morphologically similar.

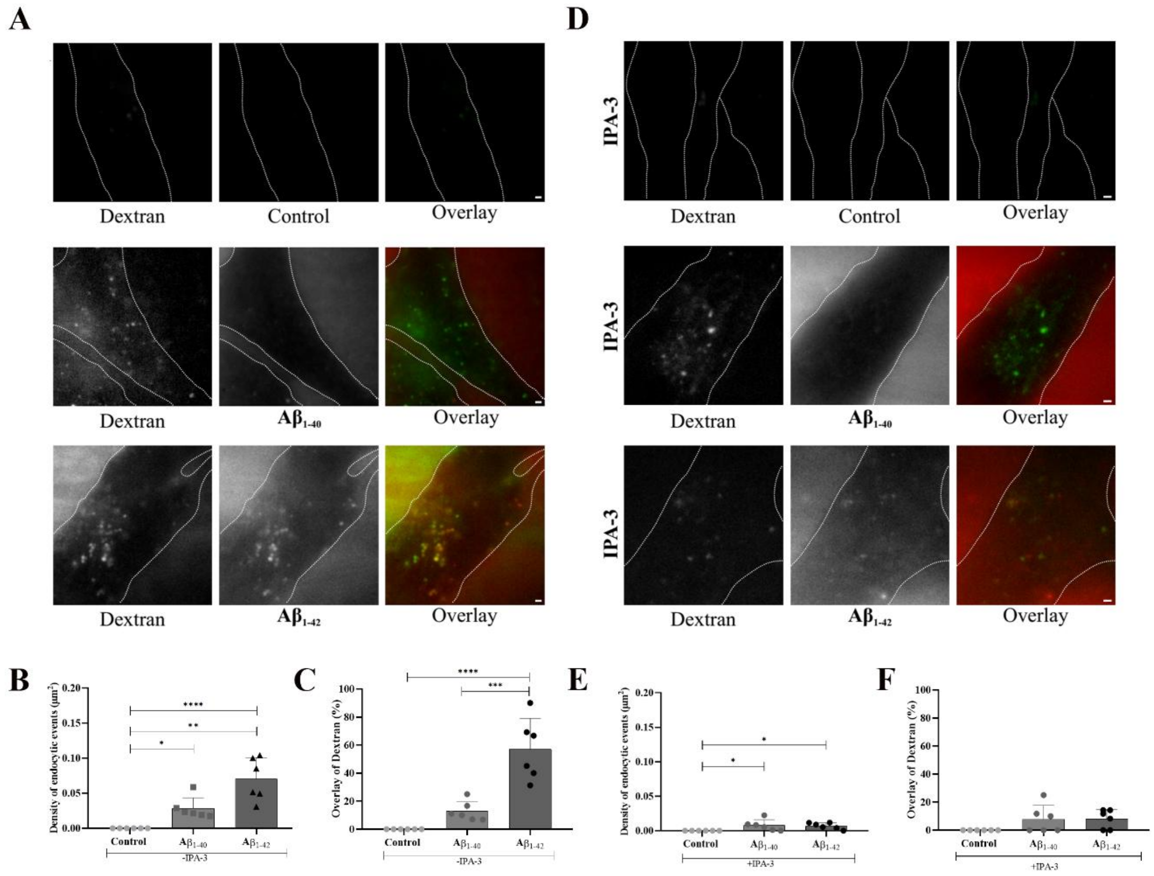


***Figure S2: Endocytosis of oAβ peptides without and with IPA-3. A****) Images showing the endocytosis of TMR-labelled o$A\beta_{1\text{-}40}$ or o$A\beta_{1\text{-}42}$ (1µM) in red channel and FITC-dextran (1 mg/ml) in the green channel, with the corresponding overlay of the images. Scale bar 1µm.* ***B)*** *Density of endocytotic events of TMR-labelled o$A\beta_{1\text{-}40}$ or o$A\beta_{1\text{-}42}$ peptides measured by counting the puncta per unit area.* ***C)*** *Percentage of colocalization of FITC-dextran with TMR-labelled o$A\beta_{1\text{-}40}$ and o$A\beta_{1\text{-}42}$.* ***D)*** *Images showing the endocytosis of TMR-labelled o$A\beta_{1\text{-}40}$ and o$A\beta_{1\text{-}42}$ in red channel and FITC-dextran in the green channel, upon IPA-3 (10 µM) treatment,*

*with the corresponding overlay of the images. Scale bar 1µm.* ***E)*** *Density of endocytotic events of TMR-labelled o*$A\beta_{1-40}$ *and o*$A\beta_{1-42}$ *upon IPA-3 treatment.* ***F)*** *Percentage of colocalization of FITC-dextran with TMR-labelled o*$A\beta_{1-40}$ *and o*$A\beta_{1-42}$ *upon IPA-3 treatment. The puncta were analyzed from 6 cells in each case, at least from 2 independent experiments. The data is presented as mean ± SD, and significance was observed with **** $p \leq 0.001$*. Statistical analysis was conducted using one-way ANOVA.*

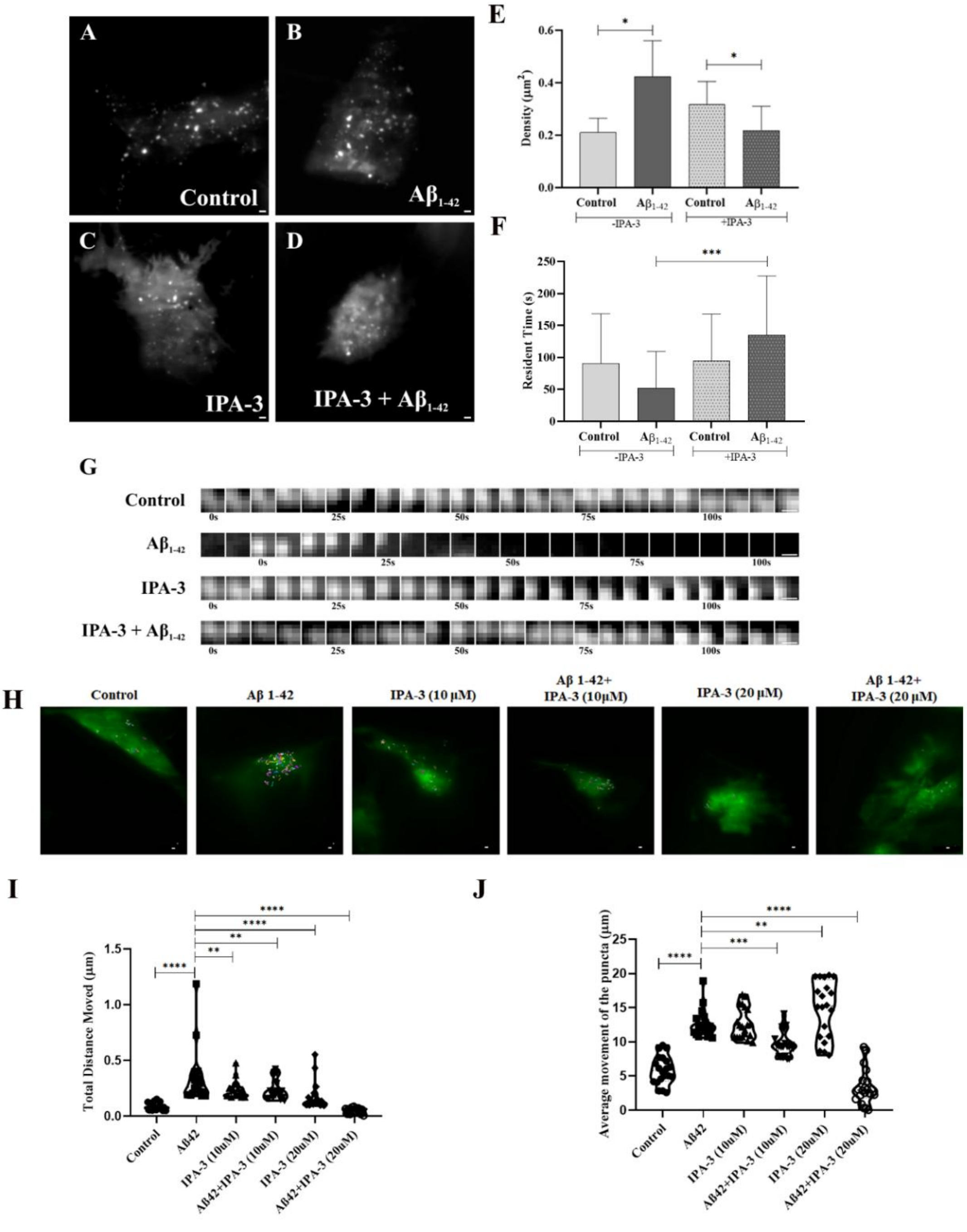

***Figure S3: Inhibition of RAB3a vesicle dynamics on the PM upon IPA-3 treatment. A-D)*** *Images of cells showing EGFP-RAB3a puncta on the PM **(A)** control **(B)** with $A\beta_{1-42}$ (1 µM) **(C)** with only IPA-3 (20 µM) and **(D)** with $A\beta_{1-42}$ upon IPA-3 (20 µM) treatment. Scale bar 1µm. **E)** Quantification of density of EGFP-RAB3a puncta on the PM without and with IPA-3 pretreatment. **F)** Residence time of RAB3a puncta on the PM upon addition of o$A\beta_{1-42}$ without or with IPA-3. **G)** Time series strip showing dynamics of RAB3a puncta on the PM without $A\beta_{1-42}$, with $A\beta_{1-42}$, with only IPA-3 treatment and, with $A\beta_{1-42}$ upon IPA-3 treatment. Scale bar 0.25µm. **H)** Image of the cell showing tracks of EGFP-RAB3a puncta on the PM of control cells, upon o$A\beta_{1-42}$ treatment, and pPAK1 inhibitor IPA-3 (at a concentration of 10 µM and 20 µM). Scale bar 1µm. **I)** Graphical representation of total distance and **J)** average distance travelled by EGFP-RAB3a puncta near the PM as shown in H. Data are presented as mean ± SEM for n=25 puncta from 5 cells in each case, at least from 2 independent experiments. The data is presented as mean ± SD, and significance was observed with *** $p \leq 0.001$. Statistical analysis was conducted using one-way ANOVA.*

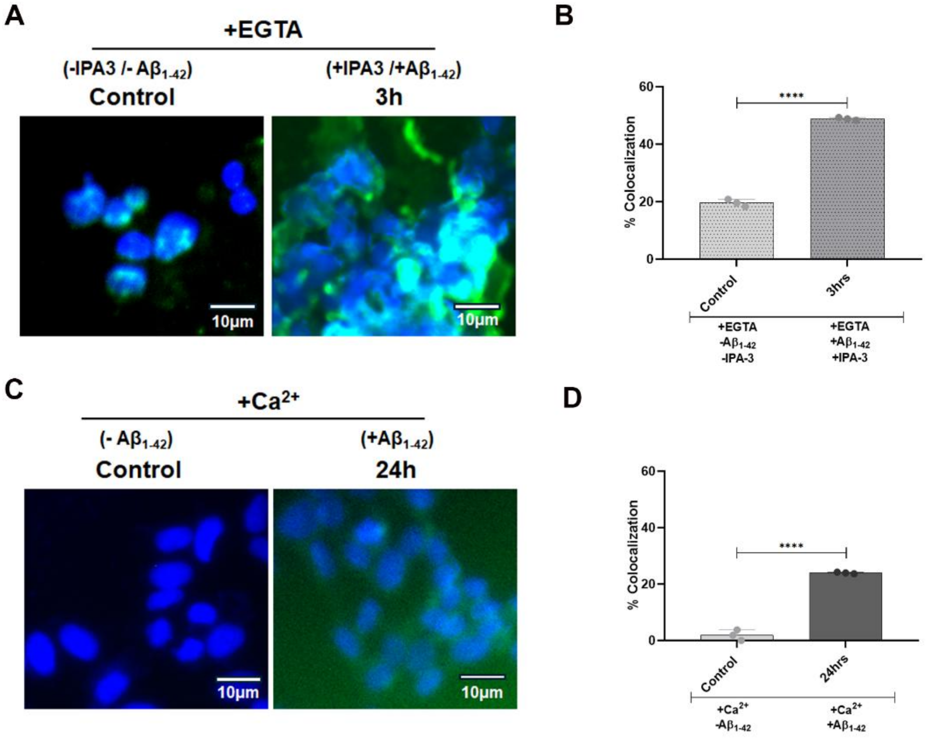


***Figure S4: Cell toxicity measured by TUNEL assay upon $Ca^{2+}$-dependent PM repair in the oAβ treated SH-SY5Y cells. A)*** *TUNEL assay images in IPA-3 (20 µM) pretreated and o$A\beta_{1-42}$ (1 µM) treated (3 hours) SH-SY5Y cells in the presence of EGTA. **B)** Quantification of the percentage of colocalization of DAPI (blue) with anti-BrdU (green) of figure (A) was*

*graphically represented.* ***C)****TUNEL assay images of oA*$\beta_{1-42}$ *treated (24 hours) cells in the presence of* $Ca^{2+}$*.* ***D)*** *The percentage of colocalization of DAPI (blue) with anti-BrdU (green) was quantified for figure (C) and graphically represented. The data is presented as mean ± SD, and significance was observed with ***** $p \leq 0.0001$*. Statistical analysis was conducted using one-way ANOVA, with a sample size of n=3.*

### Figure legends of Supplementary movies

#### Movie S1:

Time-lapse images were captured before and after addition of oAβ pepides (oA$\beta_{1-40}$ and oA$\beta_{1-42}$) at 37 ℃ using a TIRF microscope, Nikon ECLIPSE Ti2 with 100x/1.45N objective 488 excitation filter for 5 minutes with 5 seconds interval by ORCA-Flash4.0 V3 digital CMOS camera (Hamamatsu). The acquisition was through 16-bit images with 0.065 µm/pixel resolution. Dynamic distribution of EGFP-Rab3a vesicles is shown in the cells without A$\beta_{1-40}$ and A$\beta_{1-42}$ treatment in Control SH-SY5Y cells. Scale bar is 1µm.

#### Movie S2:

Time-lapse images were captured after adding oA$\beta_{1-40}$ using the TIRF microscopy at 37°C. The video was made for 5 minutes, taking the images at every 5-second interval using the CMOS camera. The video acquisition was made with 16-bit pictures at 0.065 µm/pixel resolution. The dynamic distribution of EGFP-Rab3a vesicles is shown in the cells treated with A$\beta_{1-40}$. Scale bar is 1µm.

#### Movie S3:

Time-lapse images of oA$\beta_{1-42}$ treated cells were captured using the TIRF microscopy as mentioned above. The dynamic distribution of EGFP-Rab3a vesicles was captured in the cells treated with A$\beta_{1-42}$. Faster dynamics and significantly higher distributions in the periphery were observed in the A$\beta_{1-42}$ cells compared to the control and A$\beta_{1-40}$ video. Scale bar is 1µm.
